# Paenitracins, a novel family of bacitracin-type nonribosomal peptide antibiotics produced by plant-associated *Paenibacillus* species

**DOI:** 10.1101/2025.10.21.683680

**Authors:** Nataliia V. Machushynets, Somayah S. Elsayed, Chao Du, Vladyslav Lysenko, Mercedes de la Cruz, Pilar Sanchez, Olga Genilloud, Nathaniel I. Martin, Mark R. Liles, Gilles P. van Wezel

## Abstract

The growing threat of antimicrobial resistance necessitates the discovery of novel antibiotics with activity against drug-resistant pathogens. Members of the genus *Paenibacillus* are a rich source of nonribosomal peptides (NRPs), including well-known antibiotics such as polymyxins, paenibacterin and tridecaptins. Here we use a targeted Mass-QL-based mass spectrometry approach to identify the NRPs produced by a collection of 227 taxonomically diverse plant-associated *Paenibacillus* strains, providing detailed insights into their NRP-producing potential. Using MassQL to zoom in specifically on NRPs containing basic amino acids, we discovered a novel family of bacitracins, which we designated paenitracins. The paenitracins are the first bacitracin-type peptides reported in *Paenibacillus*, and are distinguished from canonical bacitracins by three previously unseen amino acid substitutions. The paenitracins exhibit potent activity against Gram-positive pathogens, including vancomycin-resistant *Enterococcus faecium* E155. Our work provides a novel metabolomics- and genomics-guided workflow for the discovery of bioactive NRPs as a strategy to prioritize natural product chemical space and accelerate antibiotic discovery.

**IMPORTANCE:** Members of the genus *Paenibacillus* play an important role in soil ecology, producing a range of important nonribosomal peptides (NRPs) that protect their eukaryotic host. A collection of plant-associated *Paenibacillus* spp. was analyzed for their phylogenetic and metabolic diversity. We developed a novel discovery pipeline that combines feature-based molecular networking with MassQL queries to systematically prioritize bioactive NRPs containing basic amino acids. Thus we provide a comprehensive genus-wide inventory of NRPs produced by *Paenibacillus* spp. We thereby identified the paenitracins, a new subfamily of bacitracins active against multidrug-resistant Gram-positive pathogens. Our pipeline enables the discovery of novel peptidic natural products to accelerate the prioritization of chemical space for antibiotics.

## INTRODUCTION

Antimicrobial resistance (AMR) continues to challenge global health, and the discovery of new antibiotics is therefore a prerequisite to be able to keep fighting infectious diseases caused by multidrug-resistant pathogens (1). Bacterial natural products (NPs) represent a rich reservoir of antibacterial compounds, many of which have been successfully translated into clinical applications (2). A major challenge is to prioritize the huge extant genomic space for natural products (3).

Members of the bacterial phylum Bacillota (formerly Firmicutes) are well known for their potential to produce a wide range of antimicrobials, in particular nonribosomally synthesized peptides (NRPs) (4–6). Numerous bioactive NRPs produced by *Bacillus* and *Paenibacillus* have been identified, including octapeptins, paenibacterins, polypeptins, tridecaptins, etc (5–7). NRPs such as polymyxins and bacitracins that are derived from these microorganisms, have found application in clinical settings (8, 9). A large fraction of the NRPs contain positively charged amino acids and possess potent antimicrobial activities against Gram-positive and Gram-negative pathogens (10, 11). The mode of action polymyxins, involves binding to the lipid A component of lipopolysaccharides (LPSs) in the outer membrane of Gram-negative bacteria. Other NRPs, like bacitracin produced by *Bacillus subtilis* and *Bacillus licheniformis*, act as inhibitors of cell wall biosynthesis. (9) Specifically, in the presence of a divalent metal ion, most commonly Zn^2+^, bacitracin binds to and sequesters the membrane-associated phospholipid undecaprenyl pyrophosphate (C_55_PP) (12, 13). Bacitracin A is the major bioactive congener with the highest clinical relevance, and it is mainly used topically to treat infections caused by Gram-positive bacteria (14). The bacitracin family is composed of structurally similar macrocyclic dodecapeptides containing a unique N-terminal aminothiazoline moiety (15). Previously reported bacitracin A-F congeners vary at positions 1, 5, and 8 of the peptide sequence, with substitutions among Ile, Leu, and Val (16).

One of the challenges in NP discovery is the rapid identification of previously reported compounds, termed structural dereplication (17). Liquid chromatography coupled with high-resolution mass spectrometry (LC-MS) is widely used at the early stages of dereplication studies. Molecular formulas of the detected molecules are determined through the exact mass using the Seven Golden Rules, Sirius 2, and MSFINDER software; and subsequently queried in various NP databases like DNP, UNPD, ChemSpider, and REAXYS; to annotate the molecular structures (18).

In order to further identify compounds in LC-MS-based metabolite profiling workflows, the introduction of the Global Natural Products Social Molecular Networking (GNPS MN) has appeared as a powerful tool to process tandem mass spectrometry (MS) data and perform molecular networking (MN) (19). The MN concept is based on organizing and visualizing tandem MS data through a spectral similarity map, revealing the presence of homologous MS_2_ fragmentations. As structurally related compounds share similar fragmentation spectra, their nodes tend to gather together and create clusters of analogs (20). The known compounds are annotated by matching their MS^2^ spectra against an available spectral library. Moreover, combination of MN with annotation methods, such as MS2LDA (21), NAP (22), or DEREPLICATOR (23) accelerates the identification of unannotated ions (24). Mass Spec Query Language (MassQL) has been developed based on advancements in molecular networking to explore underutilized MS/MS data further (25). MassQL captures the unique characteristics of MS data, including isotopic patterns, diagnostic fragmentation, and neutral loss; and establishes a comprehensive MS terminology for searching MS patterns across datasets. Additionally, NP prioritization strategies have been applied based on the combination of molecular networking with different sources of information (e.g., genomic, taxonomic, geographic, bioactivity, and/or spectral data) (26–29). Despite the ability of current tools to dereplicate NPs upstream of the isolation process, effective strategies for the early detection of potentially bioactive NPs within crude extracts are still lacking.

In this study, we provide a full phylogenomic analysis of 227 *Paenibacillus* spp. and their potential to produce NRPs. We introduce an early-stage prioritization method that combines feature-based MN (FBMN) and MassQL search to target basic amino acid-containing NRPs with potential antimicrobial properties that are produced by a collection of plant-associated *Paenibacillus* spp. Basic amino acid signatures were detected and mapped onto a massive multi-informative molecular network. This led to the discovery of new bacitracin congeners, which we designated paenitracins, with potent bioactivity against Gram-positive pathogens, including vancomycin-resistant *E. faecium* E155.

## RESULTS AND DISCUSSION

### Taxonomic and chemical diversity of a collection of *Paenibacillus* strains

Considering the great pharmacological and biotechnological potential of the metabolites produced by *Paenibacillus*, we here analyzed 227 *Paenibacillus* isolates from the Auburn University (USA) plant-associated microbial strain collection. These *Paenibacillus* species had been isolated from the root surface of field-grown corn (*Z. mays*) and soybean (*G. max*) grown in Dunbar (Nebraska), Carrol (Iowa), Whitewater (Wisconsin), Sparta (Illinois), Troy (Ohio) and Tallassee (Alabama) (Table S1). Taxonomic analysis of the strains was conducted by comparing the 16S rRNA genes from the isolates, and a maximum-likelihood tree was constructed (Figure 1).

**Figure 1.**
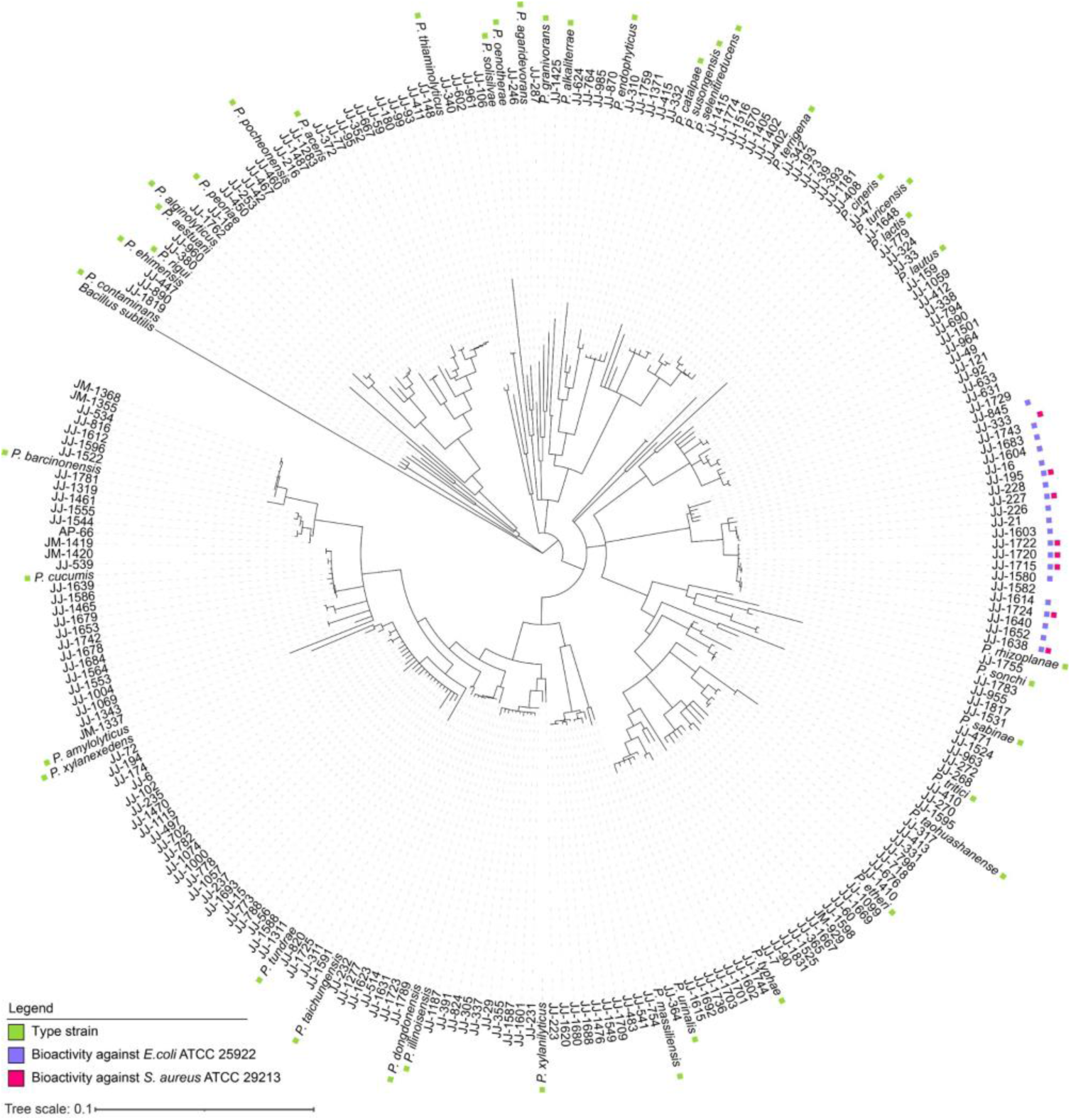
Phylogenetic tree based on 16S rRNA sequences of plant-associated *Paenibacillus* isolates and closely related type strains. The tree was obtained by neighbor-joining analysis. For an overview of the strains see Tables S1 and S2. Type strains are indicated with green squares. The tree was rooted using *Bacillus subtilis* IAM 12118 as an outgroup. Antibacterial activity of the strains against *E. coli* ATCC 25922 and *S. aureus* ATCC 29213 is indicated with purple and red squares, respectively. The tree was generated using Geneious v2024.0.3.

In search of novel NRPs, we investigated the chemical diversity of the specialized metabolites produced by *Paenibacillus* isolates. For this purpose, all 227 isolates of the collection were cultured individually on TSA media in 96 deep-well plates for 72 h and extracted with isopropyl alcohol (IPA) supplemented with 0.1% (v/v) formic acid (FA). Subsequently, crude extracts were tested against Gram-positive (*Staphylococcus aureus* ATCC 29213) and Gram-negative (*Escherichia coli* ATCC 25922) indicator strains to assess the antibacterial activity of the strains. Nine crude extracts exhibited significant growth inhibition of *S. aureus*, while 22 crude extracts showed bioactivity against *E. coli* (Figure 1, Table S1). These extracts were then subjected to LC-MS/MS analysis to identify the natural products accountable for the observed bioactivities. LC-MS/MS data were processed with MzMine 2 (30) and exported for GNPS FBMN. Prior to uploading the data to GNPS, media-derived mass features were eliminated. A mass exclusion threshold of 300 *m/z* was implemented to minimize the occurrence of mass features originating from small metabolites that are not relevant to our study. FBMN analysis resulted in generating molecular network consisting of 3096 parent ions (nodes) connected through 3692 edges (Figure 2). Characteristics of the natural products, such as annotation, *m/z* value and species-specific molecule production were visualized using Cytoscape 3.9.1 (31).

**Figure 2.**
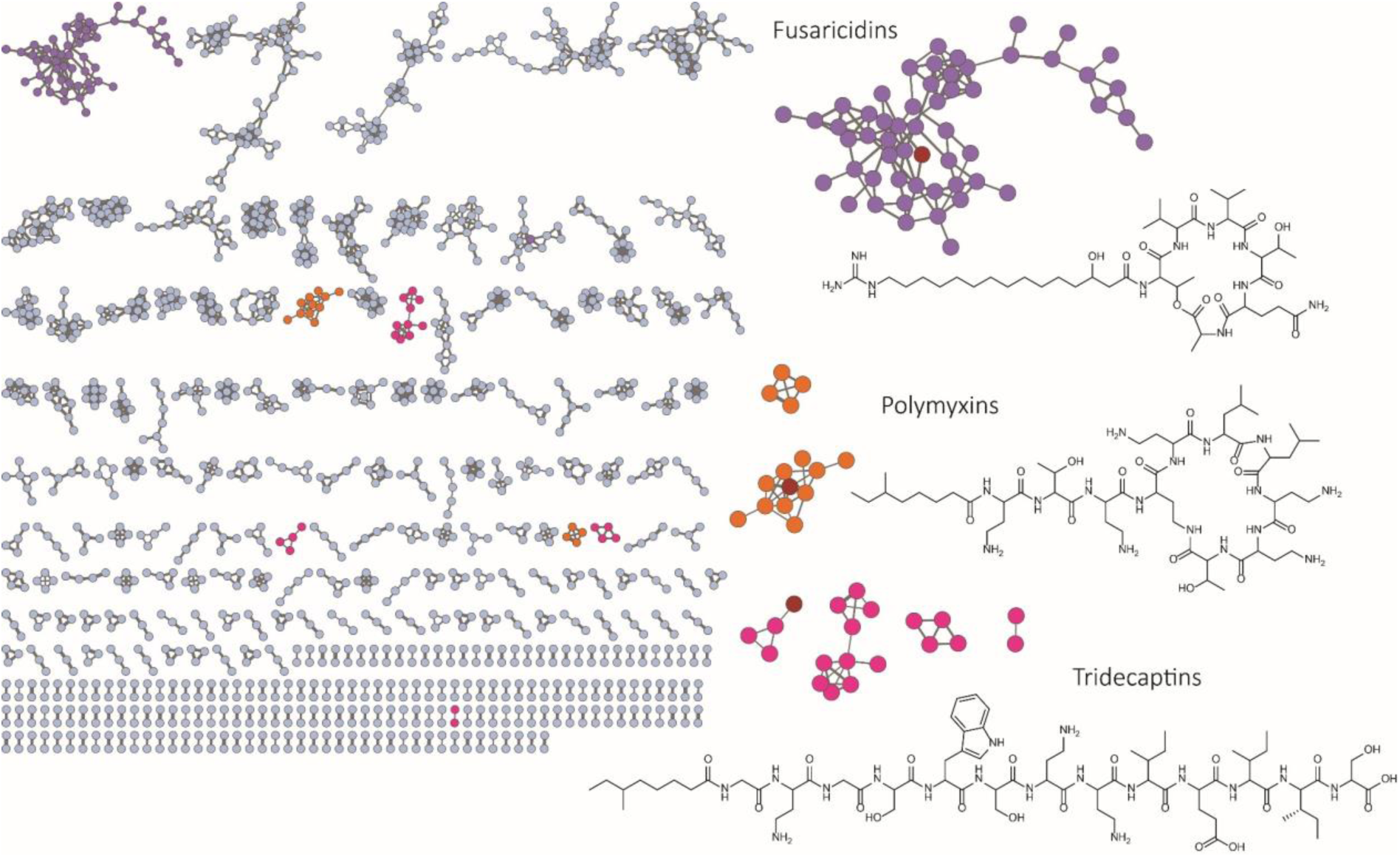
Molecular network of mass features detected in the bacterial extracts from the *Paenibacillus* isolates. Each node represents a parent ion detected in one or more strains. Different node colors represent distinct molecular families. The red nodes highlight the specific annotated compound in each family, with the corresponding chemical structure displayed next to the network.

Mass spectrometry-based molecular networking allows clustering of molecules with similar MS/MS fragmentation patterns, which stem from the similarity in their structures (19). The clusters allow visual exploration of the metabolites produced by the strains, enabling rapid dereplication of known compounds and visualization of unknown compounds (20). Known metabolites were dereplicated by matching the experimentally derived MS/MS fragmentation patterns with the annotated and curated MS/MS spectra within the GNPS database. Since the spectral library search did not work well for NRPs from *Paenibacillus*, an in-house database was created of the MS data for known lipopeptides from *Paenibacillus* and used to dereplicate known metabolites. This identified lipopeptides belonging to the families of fusaricidins, polymyxins and tridecaptins (Table S3, Figure 2, Figures S1-S3). These dereplicated mass features were used to propagate the annotation to other connected mass features. Despite the identification of a substantial number of putative congeners of known lipopeptides in the network, the majority of the produced metabolites remained unidentified.

### Targeting cationic lipopeptides using MassQL-assisted fragment search

Several of the identified lipopeptides are well-known antibiotics with potent activity, typically attributed to their net positive charge (Table S3), which enhances electrostatic interactions with the negatively charged components on the surface of bacterial cells, such as the cell membrane or lipopolysaccharides (LPSs) (32). Therefore, we sought to detect specifically those lipopeptides with potential bioactive properties that contained one or multiple positively charged amino acid residues. To accomplish this, a feature-based spectral similarity network was paired with a MassQL (33) search query to target the MS/MS spectra that contained the most common positively charged amino acids. The fragment search of *b* ions of 2,4-diaminobuturic acid (Dab) (*m/z* 101.0709), ornithine (Orn) (*m/z* 115.0871), lysine (Lys) (*m/z* 129.1027), arginine (Arg) (*m/z* 157.1089) and the immonium ion of histidine (His) (*m/z* 110.0718) was conducted with the accuracy of *m/z* 0.005 and at minimum intensity of 5% (Figure 3).

**Figure 3.**
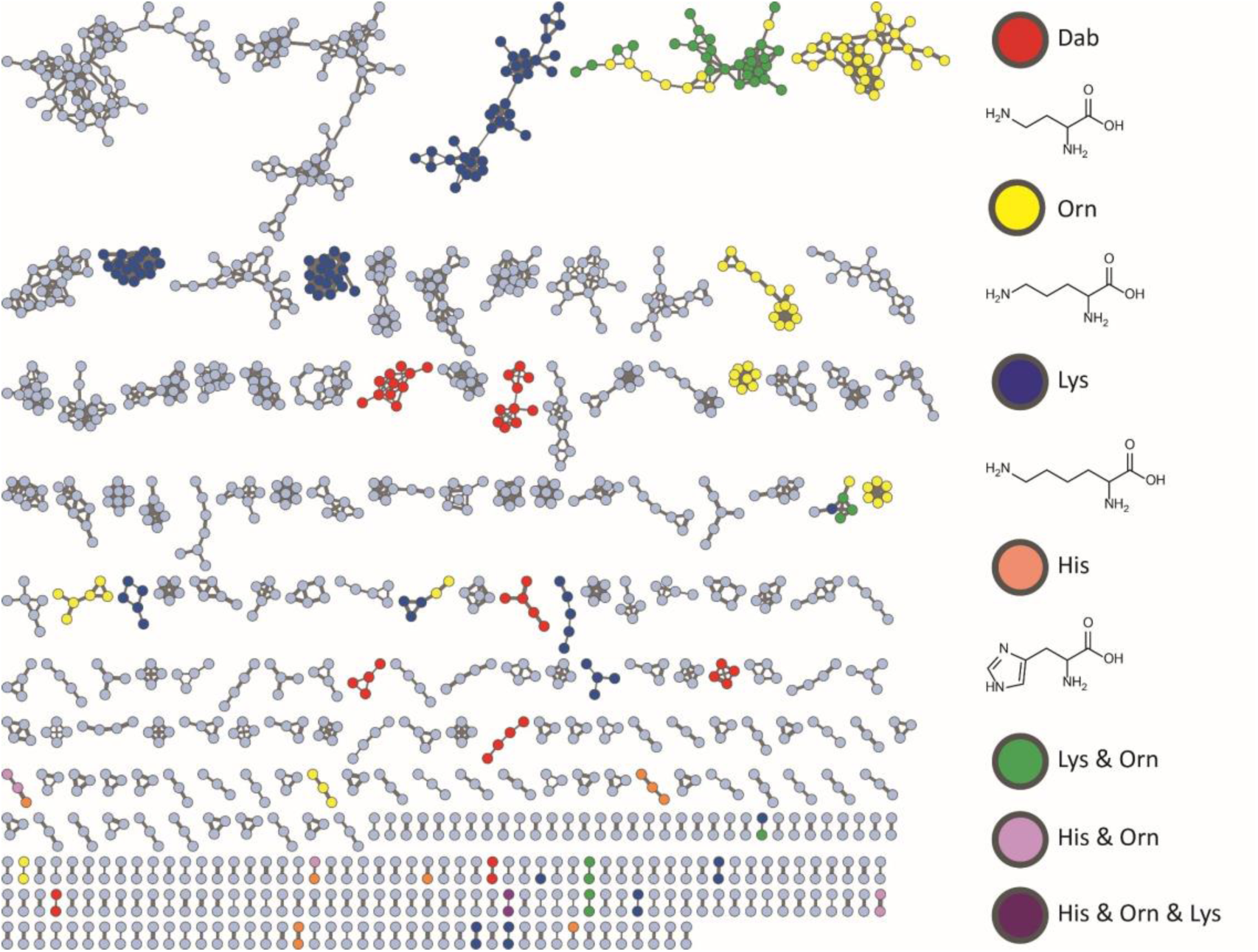
MassQL-annotated molecular network of the ions detected in the bacterial extracts from *Paenibacillus* isolates. Colors of the nodes, as indicated in the figure, highlight parent ions that contain specific basic amino acid residues (Dab, Orn, His, Lys or combinations thereof) in the structures.

MassQL search of Dab-containing parent ions highlighted previously dereplicated polymyxin and tridecaptin spectral families (Figure 4). The identification of different polymyxin (34, 35) and tridecaptin (36) congeners within numerous *Paenibacillus* extracts underscored the chemical diversity of our strain collection. Despite the structural heterogeneity of polymyxins and tridecaptins, a shared characteristic is the occurrence of multiple Dab amino acid residues. Furthermore, MassQL revealed a spectral family of compounds that contained two distinct amino acid residues, namely ornithine and lysine (Figure 5A). This cluster represented triply-charged ions with a mass range of 1500–1650 Da. Interestingly, the compounds from this cluster were produced exclusively by two isolates from the entire strain collection, namely *Paenibacillus* sp. JJ-602 and *Paenibacillus* sp. JJ-1115. These compounds could not be dereplicated by spectral library and in-house database search. Extensive literature search and manual MS/MS dereplication resulted in the annotation of these compounds as paenibacterin A congeners (Figure 5B) (10).

**Figure 4.**
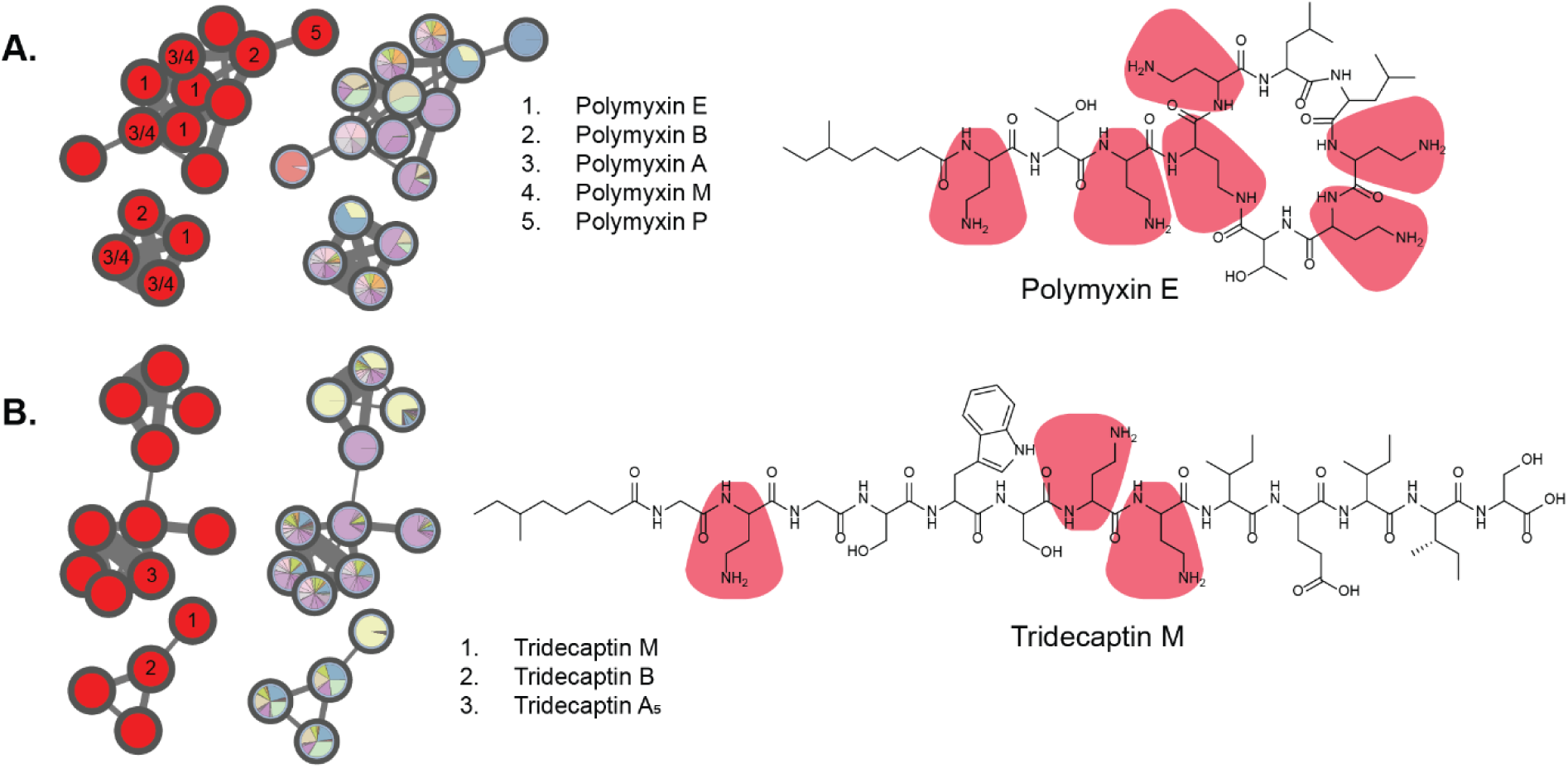
Dab-containing lipopeptide families highlighted by MassQL search. Molecular families of polymyxins (A) and tridecaptins (B) consist of multiple structural analogues, reflecting high chemical diversity. Pie charts were mapped to the nodes to represent the relative precursor ion intensities within each extract. Note that both polymyxins and tridecaptins contain multiple Dab residues.

**Figure 5.**
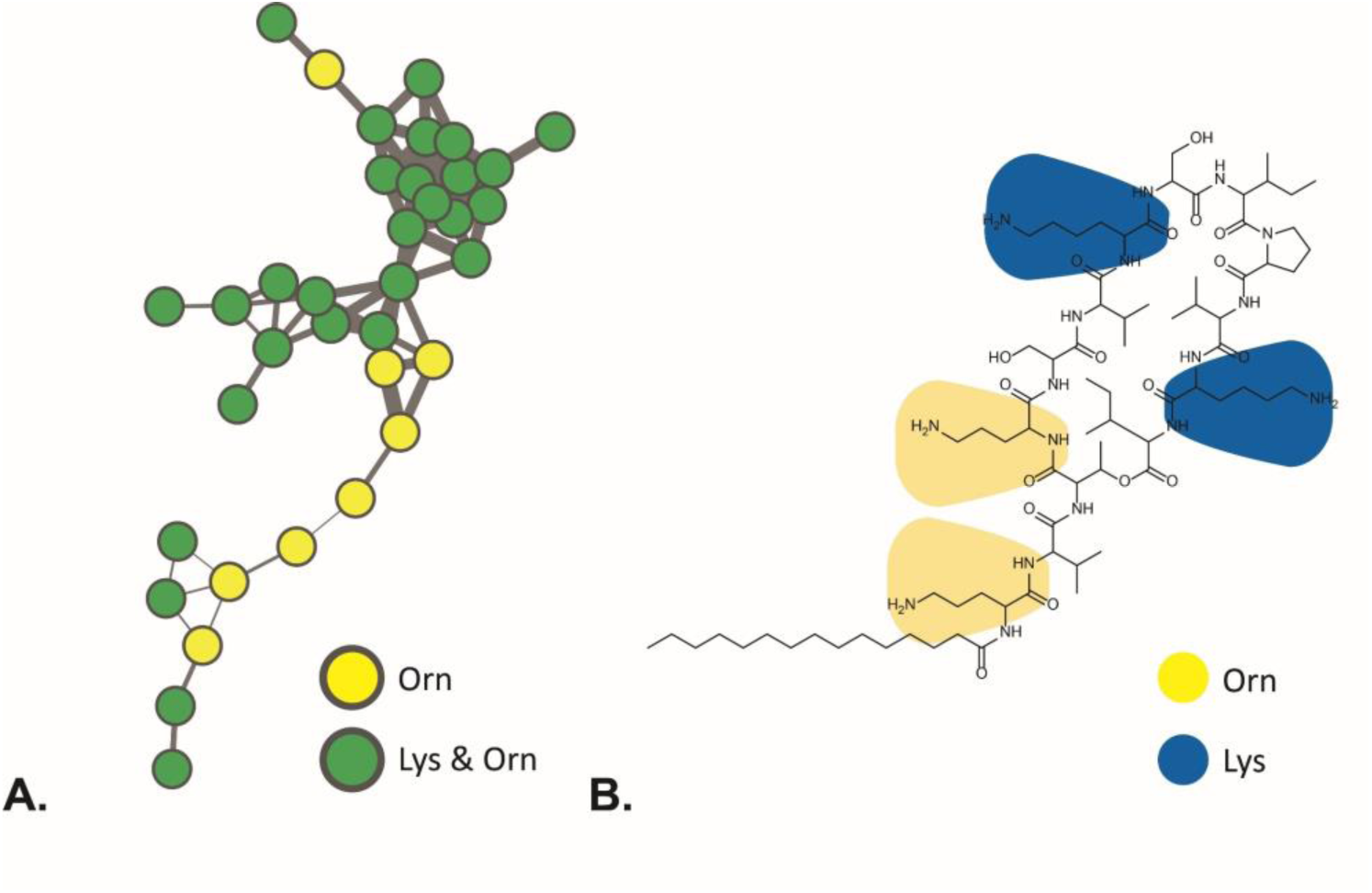
MassQL-based detection of paenibacterins. **A.** MassQL-annotated molecular family of paenibacterins. **B.** Chemical structure of paenibacterin A.

Multiple positively charged lysine and ornithine residues contribute to the overall charge of paenibacterins and enhance interactions with negatively charged LPS from the outer membrane of Gram-negative bacteria (37).

### Discovery of a novel family of bacitracins

In addition to the annotated lipopeptide classes, we identified multiple unknown molecular families that contained compounds with basic amino acids such as lysine, ornithine, and histidine (Figure 3). One molecular family of Lys-containing compounds stood out, as the production was specific to only a limited number of isolates (Figure S4). The Lys-containing molecular family could be distinguished into three sub-families (Figure 6A). Sub-family 1 and 2 represent doubly charged ions with a mass range of 1080 to 1150 Da. These compounds were similar to recently discovered lipopeptide paenilipoheptins A and B produced by *P. polymyxa* E681 (38, 39). The third sub-family comprised doubly charged molecular ions with significantly larger masses compared to paenilipoheptins (Figure 6A). The edge connecting a node in sub-family 3 (*m/z* 712.8836) with the node due to paenilipoheptin D (*m/z* 530.8307) has a cosine score of 0.6, indicating low similarity. A mirror plot comparing the MS/MS spectra of the two mass features showed no matching dipeptide fragment ions, and the only shared ions were due to a Lys residue and fragments of Trp (Figure S6).

**Figure 6.**
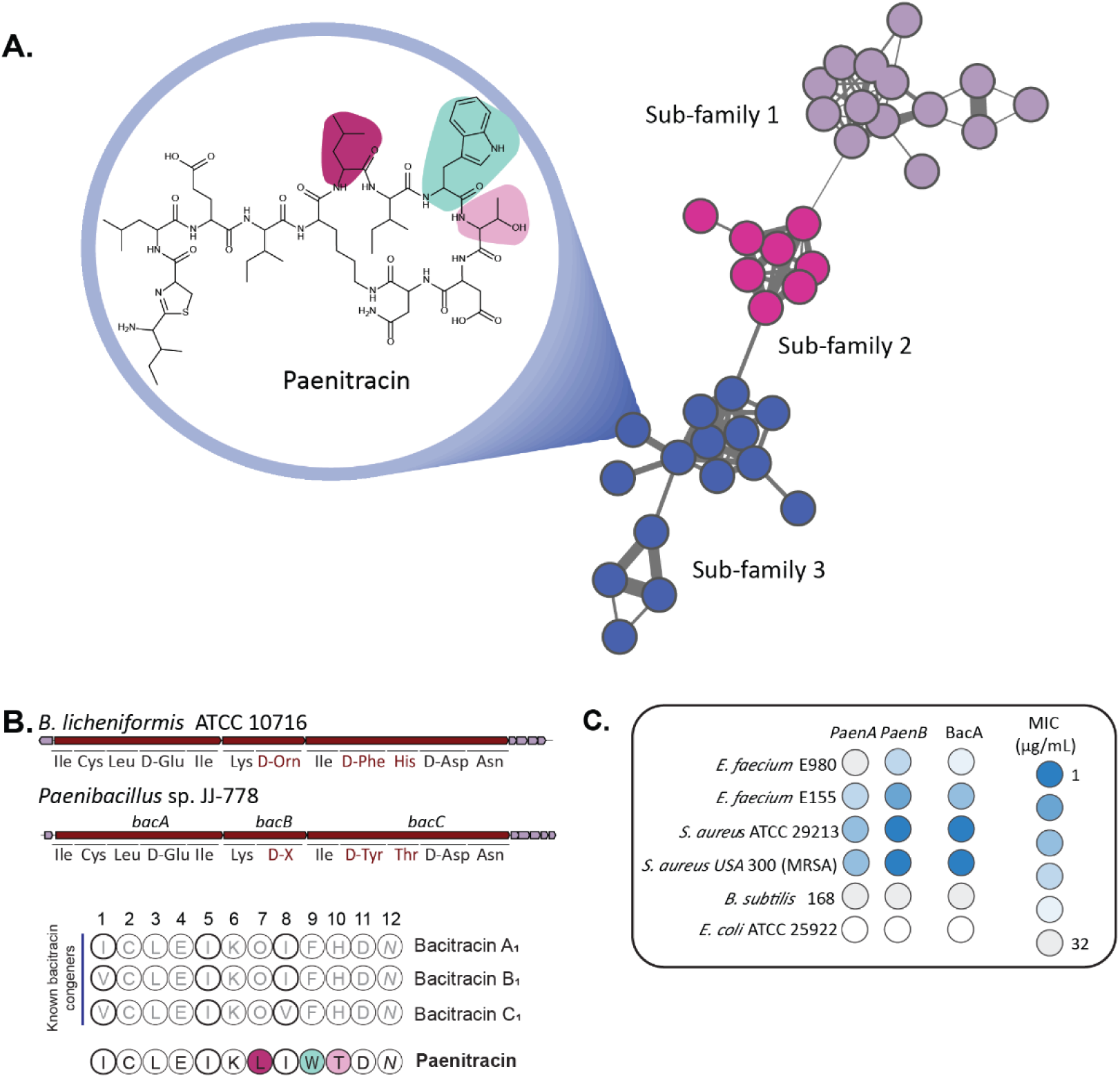
Molecular family of Lys-containing NRPs highlighted by MassQL search. **A.** Exploring the mass features from sub-family 3 led to the discovery of new bacitracin analog, designated as paenitracin. **B.** The BGC and its predicted amino acid substrates of new paenitracin from the genome of *Paenibacillus* sp. JJ-778 was compared to the reference BGC from *B. licheniformis* ATCC 10716. The amino acid sequence of paenitracin was compared to those of known bacitracins from *B. licheniformis*. Colored amino acids are unique to paenitracin. **C.** MIC data of paenitracin A and B and bacitracin against a panel of human pathogenic bacteria.

MS/MS spectra of all the mass features within this sub-family exhibited an ion with an *m/z* 199.0897. However, these mass features could not be matched to known microbial natural products. Analysis of the MS/MS spectra of the parent compound with *m/z* 712.8836 revealed that a fragment ion with an *m/z* 199.0897 has a molecular formula of C_9_H_15_N_2_OS, which corresponds to a thiazoline-containing substructure commonly found in the peptide antibiotic bacitracin, so far exclusively identified from the genus *Bacillus*. (40) Based on the fact that the *b_2_*, *b_3_, b_4_,* and *b_5_* ions were shared between the parent mass with *m/z* 712.8836 and bacitracin A (41), we concluded that their pentapeptide side chains are identical. The differences observed in *b_6_*–*b_11_* ions indicated that the structural distinctions between the two metabolites lie in the heptapeptide ring. We did not observe fragment ions corresponding to Orn, Phe or His, which are found in the structure of bacitracin A, while we could ascertain the presence of an ion corresponding to a Trp residue (*m/z* 187.0864). Moreover, the dipeptide fragment ions Trp-Leu (*m/z* 300.171), Trp-Thr (*m/z* 288.1352), and Leu-Leu (*m/z* 227.1764) were detected. By tracing the series of both *b* and *y* ions, we elucidated the amino acid sequence of the compound with *m/z* 712.8836 as (Ile-Cys)–Leu/Ile–Glu–Leu/Ile–Lys–Leu/Ile–Leu/Ile–Trp–Thr–Asp–Asn (Figure S7). The MS/MS studies used did not allow us to discriminate between Leu and Ile in positions 3, 5, 7 and 8 of the peptide amino acid sequence. Notably, this sequence diverges from any known bacitracin analogues, indicating the discovery of a novel branch of the important bacitracin-family of NRPs.

To purify the compound for more detailed studies, *Paenibacillus* sp. JJ-778 was cultured in a 10 L MHB media and subjected to extraction using Diaion® HP-20 beads, followed by elution with 100% IPA supplemented with 0.1% (v/v) formic acid. Multiple rounds of chromatographic purification resulted in the isolation of compounds (**1**) and (**2**) (Figures S8, S9). Both compounds had the same molecular formula of C_67_H_106_N_15_O_17_S, based on their exact mass of *m/z* 1424.7584 for [M+H]^+^, highly similar to the calculated mass of 1424.7611. 1D and 2D NMR analysis of **1** confirmed the amino acid sequence (Figure 7), and together with the molecular formula dictated by the accurate mass, the structure of **1** was determined as a new congener of bacitracin (Table S4, Figures S10-S16). NMR analysis identified Leu residues in positions 3 and 7, and Ile in positions 5 and 8. The ^1^H NMR spectra of **1** and **2** were almost identical, suggesting that they are diastereomers (Figure S17). Accordingly, the new compounds were designated as paenitracin A (**1**) and B (**2**).

**Figure 7.**
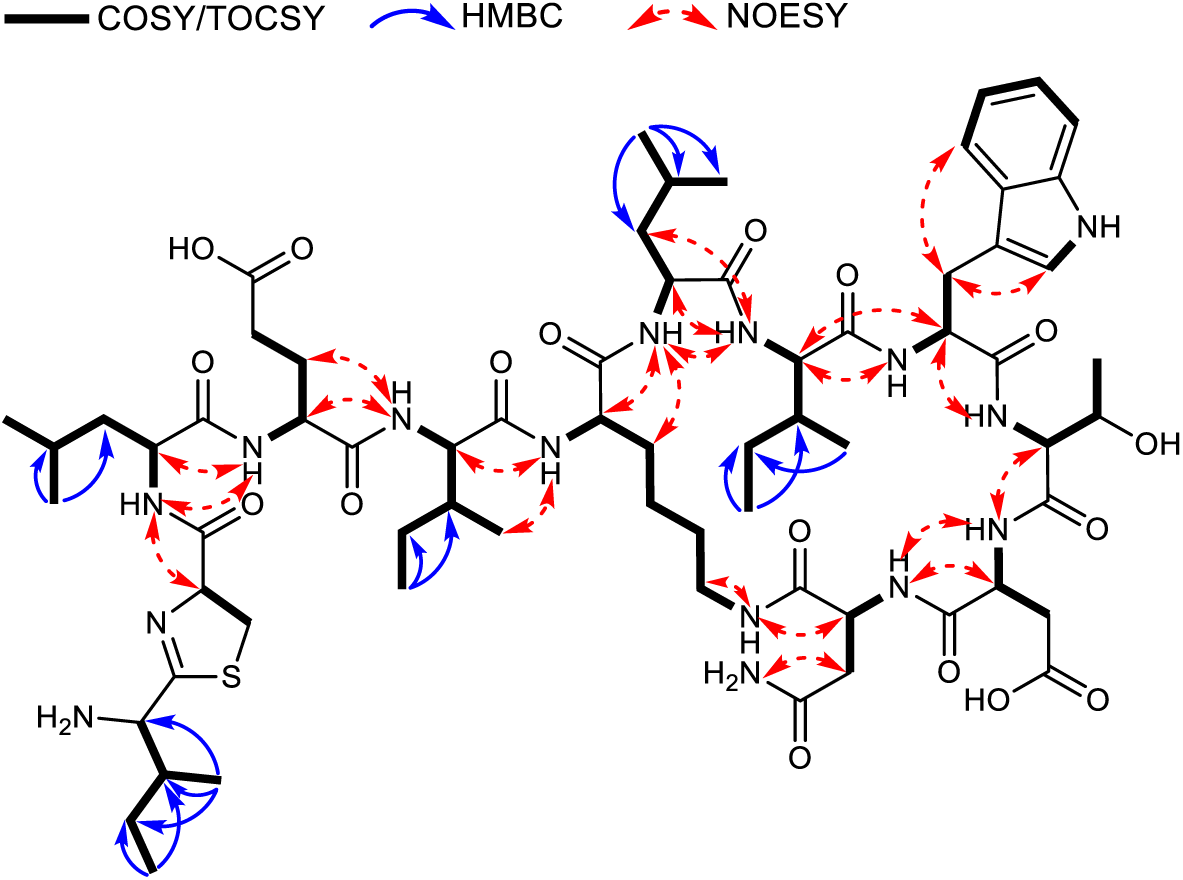
Correlations obtained by COSY, HMBC, and NOESY measurements in NMR of paenitracin A (1).

To verify the identity and stereochemistry of specific residues, compounds 1 and 2 were analyzed using Marfey’s method (Table S5, S6) (42). This analysis confirmed the presence of D-Glu, L-Lys, L-Thr, and L-Ile as single isomers. Additionally, it was found that both L and D isomers of Leu were present in the molecules, and it was also observed that the Asn and Asp residues exhibit different stereochemistry. However, the stereochemistry of the thiazoline moiety and Trp could not be determined due to their instability under the conditions used for Marfey’s analysis. The 1H NMR spectra of molecules **1** and **2** were nearly identical, with the exception of the region corresponding to the thiazoline moiety, indicating that these molecules are diastereomers (Figure S17) with stereochemical ambiguity arising from the thiazoline portion. Previous studies on bacitracin have demonstrated that racemization at the Cys position within the thiazoline moiety completely abolishes activity, whereas racemization at the Ile position only slightly impacts activity, showing a preference for L-stereochemistry (43). Consequently, we propose that the variation between molecules **1** and **2** is due to the racemization of the terminal Ile residue, with compound **1** exhibiting D-stereochemistry and compound **2** showing L-stereochemistry, as the latter was found to be marginally more potent.

Bacitracin, originally isolated from *Bacillus licheniformis* and *B. subtilis*, targets primarily Gram-positive bacteria such as *Staphylococcus*, *Streptococcus*, *Enterococcus* and *Clostridium difficile* (44). Bacitracin and its congeners are synthesized by a nonribosomal peptide synthetase (NRPS) complex encoded by the bacitracin synthetase operon, which consists of *bacA*, *bacB*, and *bacC* genes. To explore the biosynthesis of the novel paenitracin, the genome of *Paenibacillus* sp. JJ-778 was sequenced and assembled using Falcon (version 1.8.1) (45). Analysis using antiSMASH 7.1.0 (46) readily identified a 12-modular gene cluster with 77% similarity to the bacitracin BGC of *B. licheniformis* ATCC 10716. In silico analysis of the A domain substrate specificity of BacA, BacB, and BacC predicted variations in amino acid composition at positions 7, 9, and 10 compared to the well-known bacitracin A (Figure 6B).

All known bacitracin congeners isolated and identified so far exhibit similar amino acid compositions, with the Ile residues occasionally replaced by Val (Figure 6B). In contrast, paenitracin contains Leu, Trp and Thr instead of Orn, Phe and His at the amino acid positions 7, 9 and 10, respectively. This makes paenitracin a structurally distant relative of the bacitracins.

### Mode of action and bioactivity of paenitracins

Bacitracins specifically target the unique bacterial phospholipid undecaprenyl pyrophosphate (C_55_-PP). C_55_PP is the membrane-associated phospholipid that anchors lipid II and shuttles it between the cell’s cytoplasm where it is synthesized, and the membrane exterior, where it is incorporated into the growing cell wall (47). To see if paenitracin indeed has the same mode of action as bacitracin, the C_55_PP dependence of paenitracin activity was tested using antibacterial activity antagonization assays in the presence or absence of zinc (48). For this, paenitracin was mixed with C_10_-PP, a more soluble analogue of C_55_-PP, (15) and incubated with cell suspension of *S. aureus* ATCC 29213. Antagonization assays showed that the activity of paenitracin B was antagonized by C_10_PP exclusively in the presence of Zn^2+^ (Table S7). This indicates that paenitracin binds to C_55_-PP of Gram-positive bacteria in a zinc-dependent manner and thus has the same mode of actions as bacitracins.

To obtain insights into the antibacterial spectrum for paenitracin A and B, we performed minimum inhibitory concentration (MIC) assays against a panel of representative pathogenic strains, namely, *E. coli* ATCC 25922, *B. subtilis* 168, *S. aureus* ATCC 29213, *S. aureus* USA300, *E. faecium* E155 and *E. faecium* E980. The media used in the MIC assay was supplemented with 0.3 mM ZnSO_4_ to provide the divalent metal ion required for bacitracin’s activity (43). The activity of the paenitracin diastereomers was compared to that of commercially sourced bacitracin A, which served as a positive control. Paenitracin B exhibited activity against Gram-positive bacteria, with MICs ranging from 1 to 8 μg/mL, comparable to commercial bacitracin (Figure 6C). *B. subtilis* 168, previously reported to be resistant to bacitracin, also exhibited reduced susceptibility to paenitracins A and B, with MIC values of 32 μg/mL. Importantly, paenitracin B exhibited activity against vancomycin-resistant *E. faecium* E155 with MICs of 2 μg/mL, which is lower compared to the MIC of the commercially available bacitracin A of natural origin. Considering the divergent structure of paenitracin as compared to the previously reported congeners, the mode of action should be investigated further.

In conclusion, MS-based indexing of NRPs from 227 plant-associated *Paenibacillus* isolates revealed extensive chemical diversity. Molecular networking combined with MassQL identified several classes of lipopeptides, whereby we specifically focused on NRPs containing basic residues. This revealed the presence of distant relatives of bacitracin, which we designated paenitracins. Paenitracin A and B showed potent bioactivity against Gram-positive ESKAPE pathogens, including vancomycin-resistant *E. faecium* E155. Unlike previously described bacitracin variants, paenitracins show substantial structural divergence and represent the first example of bacitracin-family NRPs in *Paenibacillus*. These findings highlight the untapped chemical space within this genus and demonstrate the power of combining metabolomic indexing with targeted mining strategies to accelerate the discovery of structurally unique antibiotics.

## MATERIALS AND METHODS

### General experimental procedures

NMR spectra were recorded on a Bruker Ascend 850 MHz NMR spectrometer (Bruker BioSpin GmbH). Data were analyzed using MestReNova 14 software (Mestrelab Research, Santiago de Compostela, Spain). High-performance liquid chromatography (HPLC) purification was performed on a Waters preparative HPLC system composed of a 1525 pump, a 2707 autosampler, a 2998 PDA detector, and a Water fraction collector III. All solvents and chemicals were of HPLC or LC-MS grade, depending on the experiment.

### Isolation and identification of bacterial strains

*Paenibacillus* strains were obtained from the Auburn University Plant-Associated Microbial strain collection and had previously been isolated from the root surface of field-grown corn (*Zea mays*) and soybean (*Glycine max*) plants grown in Dunbar (Nebraska), Carrol (Iowa), Whitewater (Wisconsin), Sparta (Illinois), Troy (Ohio) and Tallassee (Alabama) (Table S1).

The phylogenetic tree was constructed using 16S rRNA gene sequences obtained through PCR amplification with universal bacterial primers 27F and 1492R (49). Each PCR amplicon was purified and used for Sanger sequencing. Subsequently, 16S rRNA gene sequences were assembled into consensus sequences and aligned with reference sequences. These include 220 sequences from our study, 41 sequences from type strains of *Paenibacillus* spp. downloaded from GenBank (Table S2) and sequence of *Bacillus subtilis* IAM 12118 (GenBank accession NR_112116.2). The alignment is done using ClustalOmega (v 1.2.2) (50). Aligned sequences are trimmed to the length of 1,452 bp to remove extending regions from a few sequences and also to preserve as many variable regions as possible. The trimmed alignment is re-aligned and a neighbor-joining consensus tree is generated using Geneious (v 2024.0.3) tree function, with Tamura-Nei genetic distance model and *B. subtilis* as outgroup. The generated tree is then uploaded to ITOL for visualization (51).

### Growth of *Paenibacillus* spp. and natural product extraction

Our method for cultivation and extraction of specialized metabolites was adapted from previously described procedures (52). Briefly, frozen stocks of 227 *Paenibacillus* spp. from the Auburn University strain collection were inoculated into 1600 µL of Tryptic Soy Agar (TSA, Bacto Soybean-Casein Digest Medium, 30 g/L) in 2.0 mL 96 deep well plates (Thermo Scientific, Nunc 2.0 mL DeepWell Plate). After 72 h, cultures were used to inoculate 200 µL Tryptic Soy Broth (TSB, Bacto Soybean-Casein Digest Medium, 30 g/L) with a pin replicator in 96 well plates. These cultures were incubated overnight at 30 °C and used as an inoculum for the second 2.0 mL 96-deep well plate containing 600 µL TSA. Plates were sealed with 96 Well-Cap Mats (Thermo Scientific, Nunc 96 Well-Cap Mats), and incubated at 30 °C for 72 h. The cultures were extracted with 300 µL 100% isopropanol acidified with 0.1% formic acid. The plates were resealed with the same 96 Well-Cap Mats, sonicated for 10 min, and extracted for an additional 50 min. 200 µL of these crude extracts were transferred into a pre-washed 96 well plate (Agilent Technologies, 96 well plates, 0.5 mL, polypropylene) and lyophilized to dryness. The extraction protocol was repeated once more for a total extract volume of 400 µL. Dried samples were redissolved in 160 µL of methanol:water (1:1 v/v).

### Genome sequencing, assembly and annotation

*Paenibacillu*s sp. JJ-778 was grown in TSB at 30 °C and 220 rpm for 24 h. DNA was extracted as described (53). DNA quality was verified by agarose gel electrophoresis. PacBio sequencing and assembly was performed by Novogene (UK). Generally, libraries were prepared using SMRTbell template prep kit (PacBio, USA) according to manufacturer instructions. Sequencing was then performed using PacBio Sequel platform in continuous long reads mode. Assembly was done using Falcon (version 1.8.1) (45). The complete genome sequence of *Paenibacillus* sp. JJ-778 has been deposited in GenBank under the accession number CP194855. BGCs in these genomes were annotated using AntiSMASH (version 7.1.0) (46).

### Data-dependent LC-ESI-HRMS/MS

LC-MS/MS acquisition was performed using Shimadzu Nexera X2 ultra-high-performance liquid chromatography (UPLC) system, with an attached photodiode array detector (PDA), coupled to Shimadzu 9030 QTOF mass spectrometer, equipped with a standard electrospray ionization (ESI) source unit. A total of 2 µL was injected into a Waters Acquity HSS C_18_ column (1.8 μm, 100 Å, 2.1 × 100 mm) and data acquisition was performed as previously described (54). Briefly, the gradient used was 5% B for 1 min, 5–85% B for 9 min, 85–100% B for 1 min, and 100% B for 4 min. The column was re-equilibrated to 5% B for 3 min before the next run was started. All the samples were analyzed in positive polarity, using data-dependent acquisition mode. In this regard, full scan MS spectra (*m/z* 100–1700, scan rate 10 Hz, ID enabled) were followed by two data-dependent MS/MS spectra (*m/z* 100–1700, scan rate 10 Hz, ID disabled) for the two most intense ions per scan. The ions were fragmented using collision-induced dissociation (CID) with fixed collision energy (CE 20 eV) and excluded for 1 s before being re-selected for fragmentation. The parameters used for the ESI source were: interface voltage 4 kV, interface temperature 300 °C, nebulizing gas flow 3 L/min, and drying gas flow 10 L/min.

### MZmine 2 parameters

Prior to statistical analysis, mzXML files were imported into Mzmine 2.53 (30) and processed as previously described (54). Briefly, mass ion peaks were detected for MS^1^ and MS^2^ at a noise level of 2.0E2 and 0.0E0, respectively (positive polarity, mass detector: centroid), and their chromatograms were built using ADAP chromatogram builder. The detected peaks were smoothed, and the chromatograms were deconvoluted (algorithm: local minimum search; chromatographic threshold: 90; search minimum in RT range: 0.05; minimum relative height: 1%; minimum ratio of peak top/edge: 2; peak duration: 0.03–3.00 min). The detected peaks were deisotoped (monotonic shape; maximum charge: 3; representative isotope: most intense). Peak lists from different samples were aligned (weight for RT = weight for *m/z* = 50; compare the isotopic pattern with a minimum score of 50%). Missing peaks detected in at least one of the samples were filled with the gap-filling algorithm (RT tolerance: 0.1 min*, m/z* tolerance was set to 0.002 *m/z* or 15 ppm). Duplicate peaks were filtered. Only the features with MS/MS data were exported to a GNPS-FBMN.

In addition, all features that originate from the culture medium were removed by retaining only features with an average peak intensity of at least 100 times greater in the bacterial extracts than in the culture medium extracts. Features that are present in the blanks with an intensity higher than 5000 were also removed. In the end, features with *m/z* less than 300 were filtered out.

### Molecular network analysis and MassQL search

The resulting feature quantification table (CSV file) and MS/MS spectrum files (in mgf format) were uploaded to the GNPS webserver (http://gnps.ucsd.edu) (19). The LC-MS/MS data for the 227 *Paenibacillus* isolates were analyzed using the Feature-Based Molecular Networking tool (FBMN) (20). Briefly, the precursor ion mass tolerance was set to 0.005 Da and the MS/MS fragment ion tolerance to 0.05 Da. A molecular network was then created where edges were filtered to have a cosine score above 0.5 and more than 3 matched peaks. The molecular networks were visualized using Cytoscape software version 3.9.1 (31) and displayed using an unweighted force-directed layout. The data are publicly accessible in the MassIVE repository (MSV000094386) and the networking results and parameters can be found at https://gnps.ucsd.edu/ProteoSAFe/status.jsp?task=2f1e0ae1728142249ac4e841d5a72ef4. The MassQL tool version 31.4 (33) was employed to search for specific MS/MS fragments that correspond to the *b* or immonium ions of positively charged amino acids. The fragment search of *b* ions of 2,4-diaminobuturic acid (*m/z* 101.0709), ornithine (*m/z* 115.0871), lysine (*m/z* 129.1027), and arginine (*m/z* 157.1089); together with the immonium ion of histidine (*m/z* 110.0718), was conducted with an accuracy of *m/z* 0.005 and minimum intensity of 5%.

### Up-scale fermentation, extraction, and isolation

*Paenibacillus* sp. JJ-778 was grown at 30 °C on tryptic soy agar (TSA) for 72 h and three colonies were inoculated into TSB and incubated at 30 °C overnight. This inoculum (1%) was used to inoculate thirteen 2 L Erlenmeyer flasks containing 0.75 L of sterile MHB and fermented at 30 °C while shaking at 200 rpm for 72 h. To extract the specialized metabolites produced by *Paenibacillus* sp. JJ-778 supernatant was collected by centrifugation and subjected to extraction using Diaion® HP-20 (Resindion, Mitsubishi), followed by elution with 100% IPA supplemented with 0.1% (v/v) formic acid. The crude extract was adsorbed onto silica gel (pore size 60 Å, 70–230 mesh, Sigma Aldrich), and loaded on a silica column, which was eluted using varying gradients of EtOAc/MeOH. The fractions eluted with EtOAc-MeOH (1:3) supplemented with 0.1% (v/v) formic acid were combined, reconstituted in 50% ACN and injected into the preparative SunFire column (19 × 150 mm), which was eluted with H_2_O:ACN gradient of 15–60% in 30 min, at a flow rate of 15 mL/min, to yield three fractions. Fraction 3 was further separated on the semi-preparative SunFire column (10 × 250 mm) at 3 mL/min using a H_2_O–ACN gradient of 30– 50% in 30 min to yield paenitracin A (**1**, 0.3 mg) and paenitracin B (**2**, 0.2 mg).

### Marfey’s analysis

The stereochemistry of chiral centers present at α carbons were assigned by applying derivatization methods coupled with chromatographic analysis. The advanced Marfey’s method using L-FDAA (1-fluoro-2-4-dinitrophenyl-5-L-alanine amide) established the absolute configurations of amino acids (55). The general method for Marfey’s analysis was conducted as described) (42). Briefly, a sample of peptide (30 µg) in 6M HCl (100 µL) was heated to 100 °C in a sealed vial for 8−12 h using heating block, after which the hydrolysate was concentrated to dryness at 40 °C under a stream of dry N_2_. The hydrolysate was then treated with 1 M NaHCO_3_ (20 µL) and L-FDAA (1% solution in acetone, 40 µL) at 40 °C for 1 h, after which the reaction was neutralized with 1 M HCl (20 µL). An aliquot of the analyte was diluted 50 times with H_2_O/ACN (1:1) and injected (2 µL) into an HRMS instrument following the standard protocol of the analysis. The analyte amino acid content was assessed by comparison to authentic standards. The authentic standards were prepared via a similar procedure.

### Antibacterial activity assays and MIC determination

For *E. coli* ATCC 25922 and *S. aureus* ATCC 29213, single colonies were inoculated into 20 mL of MHII broth medium in 250 mL Erlenmeyer flask and incubated overnight at 37 °C with shaking at 220 rpm and then diluted to obtain an assay inoculum of 5 x 10^5^ CFU/mL. Aztreonam and vancomycin were used as reference positive controls for each pathogen. Absorbance at OD_612_ nm was measured at *T*_0_ (zero time) using an EnVision™ Microplate Reader (PerkinElmer), and assay plates were incubated at 37 °C. After that, absorbance at OD_612_ nm was measured at *T_f_* (final time) using the same microplate reader equipment. The percentage of growth inhibition was calculated using the following normalization:

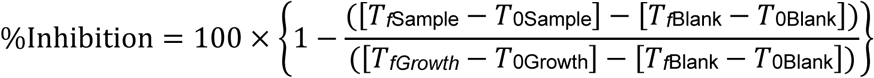

*T*_0Sample_, the absorbance of the strain growth in the presence of sample measured at zero time;
*T_f_*_Sample_, the absorbance of the strain growth in the presence of sample measured at final time;
*T*_0Growth_, the absorbance of the strain growth in the absence of sample measured at zero time;
*T_f_*_Growth_, the absorbance of the strain growth in the absence of sample measured at the final time;
*T*_0Blank_, the absorbance of the broth medium (blank) measured at zero time;
*T_f_*_Blank_, the absorbance of the broth medium (blank) measured at the final time.

The Genedata Screener software (Genedata, Inc., Basel, Switzerland) was used to process and analyze the data. The activity of the samples was expressed as a percentage of growth inhibition, where (−100%) represented the total growth inhibition of the target microorganism and 0% represented the total growth of the target microorganism in the assay. The Z’ factor predicts the robustness of an assay by considering the mean and standard deviation of both positive and negative controls (56). The robust Z’ factor (RZ’ factor) is based on the Z’ factor, but standard deviations and means are replaced by the robust standard deviations and medians, respectively. In all experiments performed in this work, the RZ’ factor obtained was between 0.9173 and 0.9684.

For MIC tests, blood agar plates were inoculated with glycerol stocks of *E. coli* ATCC 25922, *B. subtilis* 168, *S. aureus* ATCC 29213, *S. aureus* USA300 (clinical isolate from Texas Children’s Hospital, USA), *E. faecium* E155 and *E. faecium* E980 and incubated overnight at 37 °C. Following incubation, 3 mL of tryptic soy broth (TSB) was inoculated with an individual colony and incubated at 37 °C with shaking at 220 rpm until the cultures reached the exponential phase. The bacterial suspensions were then diluted 100-fold in cation-adjusted Mueller-Hinton broth (MHB) or Luria-Bertani (LB) broth supplemented with 0.3 M ZnSO_4_ to reach a bacterial cell density of 10^6^ CFU/mL. In parallel, test compounds were 2-fold serially diluted with cation-adjusted MHB or LB supplemented with 0.3 M ZnSO_4_ in polypropylene 96-well plates to achieve a final volume of 50 μL per well. An equal volume of bacterial suspension (10^6^ CFU/mL) was added to the wells. The well plates were sealed with an adhesive membrane and incubated for 20 hours at 37 °C. The reported MIC values are based on three technical triplicates and are defined as the lowest concentration of the compound that prevented the visible growth of bacteria.

### In vitro antagonization of paenitracin antibacterial activity

Antibacterial activity antagonization assay was performed as previously described (48). Briefly, geranyl pyrophosphate ammonium salt (C_10_PP) (1 mg/mL in methanol [aqueous 10 mM NH_4_OH (7:3), Sigma Aldrich, Taufkirchen, Germany] was used as an antagonist of paenitracin A in either the presence or absence of 0.3 mM ZnSO_4_. Paenitracin A was mixed with C_10_PP (five-fold molar excess relative to paenitracin) or added to empty wells (absence of antagonist) giving in all cases 50 μL of total volume in each well with paenitracin A at 4× MIC in the presence and absence of antagonist as well as in the presence and absence of 0.3 mM ZnSO_4_, in triplicate. Subsequently, 50 μL of *S. aureus* ATCC 29213 with OD_600_ of 0.005 was added to the test compounds in the microtiter plate to achieve a final antibiotic concentration of 2× MIC. The samples were incubated at 37 °C for 24 h with constant shaking and inspected for visible bacterial growth.

## Acknowledgments

We thank D. van der Horst for technical assistance with LC-MS analysis, to Jesús Martín for assistance in compound annotation and to Marnix Medema for stimulating discussions. The work was supported by the NACTAR program of The Netherlands Organization for Scientific Research (NWO), Grant 16440.

