## Supplemental Information for "Paenitracins, a novel family of bacitracin-type nonribosomal peptide antibiotics produced by plant-associated *Paenibacillus* species"

[c] Fundación MEDINA, Health Sciences Technology Park, Avda Conocimiento 34, 18016, Granada, Spain.

[d] Department of Biological Sciences, Auburn University, 120 W Samford Av, AL 36849, Auburn, AL, USA

[e] Department of Microbial Ecology, Netherlands Institute of Ecology, Droevendaalsesteeg 10, 6708 PB, Wageningen, The Netherlands

**Table S1. Overview of 227 *Paenibacillus* spp. from the Auburn University Plant-Associated Microbial strain collection.**

| Isolate | Top hit | Pairwise similarity (%) | Crop | Rhizoplane/ Endophyte | Root # | City | State | Bioactivity against *E.coli* ATCC 25922 | Bioactivity against *S. aureus* ATCC 29213 |
|---|---|---|---|---|---|---|---|---|---|
| **JJ-6** | *P. xylanexedens* | 99.39 | Corn | Endophyte | 1 | Dunbar | Nebraska | – | – |
| **JJ-7** | *P. typhae* | 98.70 | Corn | Endophyte | 1 | Dunbar | Nebraska | – | – |
| **JJ-15** | *P. amylolyticus* | 99.59 | Corn | Endophyte | 3 | Dunbar | Nebraska | – | – |
| **JJ-16** | *P. peoriae* | 99.59 | Corn | Endophyte | 3 | Dunbar | Nebraska | + | – |
| **JJ-18*** | *P. alginolyticus* | 99.22 | Corn | Endophyte | 3 | Dunbar | Nebraska | – | – |
| **JJ-21*** | *P. peoriae* | 99.66 | Corn | Endophyte | 3 | Dunbar | Nebraska | + | – |
| **JJ-29** | *P. illinoisensis* | 99.65 | Corn | Endophyte | 4 | Dunbar | Nebraska | – | – |
| **JJ-33** | *P. lautus* | 99.37 | Corn | Endophyte | 4 | Dunbar | Nebraska | – | – |
| **JJ-39** | *P. terrigena* | 99.25 | Corn | Endophyte | 6 | Dunbar | Nebraska | – | – |
| **JJ-42** | *P. pectinilyticus* | 98.78 | Corn | Endophyte | 6 | Dunbar | Nebraska | – | – |
| **JJ-47** | *P. cineris* | 100.00 | Corn | Endophyte | 7 | Dunbar | Nebraska | – | – |
| **JJ-49** | *P. lautus* | 99.16 | Corn | Endophyte | 7 | Dunbar | Nebraska | – | – |
| **JJ-56** | *P. amylolyticus* | 99.45 | Corn | Endophyte | 9 | Dunbar | Nebraska | – | – |
| **JJ-59** | *P. aceris* | 98.61 | Corn | Endophyte | 9 | Dunbar | Nebraska | – | – |
| **JJ-60** | *P. etheri* | 99.65 | Corn | Endophyte | 9 | Dunbar | Nebraska | – | – |
| **JJ-72** | *P. amylolyticus* | 99.80 | Corn | Endophyte | 12 | Dunbar | Nebraska | – | – |
| **JJ-73** | *P. terrigena* | 99.04 | Corn | Endophyte | 12 | Dunbar | Nebraska | – | – |
| **JJ-77** | *P. aceris* | 99.65 | Corn | Endophyte | 12 | Dunbar | Nebraska | – | – |
| **JJ-90** | *P. typhae* | 98.70 | Corn | Endophyte | 14 | Dunbar | Nebraska | – | – |
| **JJ-92** | *P. lautus* | 99.30 | Corn | Endophyte | 14 | Dunbar | Nebraska | – | – |
| **JJ-93** | *P. aceris* | 98.75 | Corn | Endophyte | 14 | Dunbar | Nebraska | – | – |
| **JJ-95** | *P. aceris* | 99.29 | Corn | Endophyte | 15 | Dunbar | Nebraska | – | – |
| **JJ-99** | *P. aceris* | 98.75 | Corn | Endophyte | 15 | Dunbar | Nebraska | – | – |
| **JJ-102** | *P. xylanexedens* | 99.39 | Corn | Endophyte | 16 | Dunbar | Nebraska | – | – |
| **JJ-106** | *P. marinesediminis* | 99.59 | Soybean | Endophyte | 1 | Dunbar | Nebraska | – | – |
| **JJ-121** | *P. lautus* | 99.23 | Soybean | Endophyte | 5 | Dunbar | Nebraska | – | – |
| **JJ-148** | *P. aceris* | 98.68 | Soybean | Endophyte | 11 | Dunbar | Nebraska | – | – |
| **JJ-159** | *P. lautus* | 99.23 | Soybean | Endophyte | 13 | Dunbar | Nebraska | – | – |
| **JJ-174** | *P. amylolyticus* | 99.25 | Soybean | Endophyte | 15 | Dunbar | Nebraska | – | – |
| **JJ-180** | *P. aceris* | 98.75 | Soybean | Endophyte | 16 | Dunbar | Nebraska | – | – |
| **JJ-193** | *P. terrigena* | 99.52 | Corn | Endophyte | 1 | Dunbar | Nebraska | – | – |
| **JJ-194** | *P. amylolyticus* | 99.32 | Corn | Endophyte | 3 | Dunbar | Nebraska | – | – |
| **JJ-195** | *P. peoriae* | 99.72 | Corn | Endophyte | 3 | Dunbar | Nebraska | + | + |
| **JJ-216** | *P. pocheonensis* | 99.04 | Corn | Rhizoplane | 1 | Dunbar | Nebraska | – | – |
| **JJ-223** | *P. xylanilyticus* | 98.37 | Corn | Rhizoplane | 1 | Dunbar | Nebraska | – | – |
| **JJ-226** | *P. peoriae* | 99.79 | Corn | Rhizoplane | 2 | Dunbar | Nebraska | + | – |
| **JJ-227** | *P. peoriae* | 99.79 | Corn | Rhizoplane | 2 | Dunbar | Nebraska | + | + |
| **JJ-228** | *P. peoriae* | 99.79 | Corn | Rhizoplane | 2 | Dunbar | Nebraska | + | – |
| **JJ-231** | *P. illinoisensis* | 99.86 | Corn | Rhizoplane | 3 | Dunbar | Nebraska | – | – |
| **JJ-232** | *P. dongdonensis* | 99.19 | Corn | Rhizoplane | 3 | Dunbar | Nebraska | – | – |

| | | | | | | | | | |
|---|---|---|---|---|---|---|---|---|---|
| **JJ-235** | *P. amylolyticus* | 99.66 | Corn | Rhizoplane | 3 | Dunbar | Nebraska | – | – |
| **JJ-237** | *P. amylolyticus* | 99.66 | Corn | Rhizoplane | 3 | Dunbar | Nebraska | – | – |
| **JJ-246** | *P. oenotherae* | 98.63 | Corn | Rhizoplane | 5 | Dunbar | Nebraska | – | – |
| **JJ-253** | *P. pectinilyticus* | 98.78 | Corn | Rhizoplane | 6 | Dunbar | Nebraska | – | – |
| **JJ-268** | *P. tritici* | 99.73 | Corn | Rhizoplane | 9 | Dunbar | Nebraska | – | – |
| **JJ-270** | *P. tritici* | 100.00 | Corn | Rhizoplane | 9 | Dunbar | Nebraska | – | – |
| **JJ-272** | *P. tritici* | 98.90 | Corn | Rhizoplane | 9 | Dunbar | Nebraska | – | – |
| **JJ-287** | *P. agaridevorans* | 97.41 | Corn | Rhizoplane | 13 | Dunbar | Nebraska | – | – |
| **JJ-305** | *P. illinoisensis* | 99.72 | Corn | Rhizoplane | 16 | Dunbar | Nebraska | – | – |
| **JJ-310** | *P. endophyticus* | 99.52 | Soybean | Rhizoplane | 1 | Dunbar | Nebraska | – | – |
| **JJ-311** | *P. amylolyticus* | 98.64 | Soybean | Rhizoplane | 1 | Dunbar | Nebraska | – | – |
| **JJ-317** | *P. 'taohuashanense'* | 99.86 | Soybean | Rhizoplane | 1 | Dunbar | Nebraska | – | – |
| **JJ-324** | *P. lautus* | 99.37 | Soybean | Rhizoplane | 2 | Dunbar | Nebraska | – | – |
| **JJ-329** | *P. 'yonginensis'* | 97.47 | Soybean | Rhizoplane | 3 | Dunbar | Nebraska | – | – |
| **JJ-331** | *P. 'taohuashanense'* | 98.66 | Soybean | Rhizoplane | 3 | Dunbar | Nebraska | – | – |
| **JJ-332** | *P. catalpae* | 99.59 | Soybean | Rhizoplane | 3 | Dunbar | Nebraska | – | – |
| **JJ-333** | *P. jamilae* | 99.86 | Soybean | Rhizoplane | 3 | Dunbar | Nebraska | – | + |
| **JJ-337** | *P. illinoisensis* | 99.79 | Soybean | Rhizoplane | 4 | Dunbar | Nebraska | – | – |
| **JJ-338** | *P. lautus* | 99.37 | Soybean | Rhizoplane | 4 | Dunbar | Nebraska | – | – |
| **JJ-340** | *P. thiaminolyticus* | 99.79 | Soybean | Rhizoplane | 4 | Dunbar | Nebraska | – | – |
| **JJ-342** | *P. terrigena* | 99.52 | Soybean | Rhizoplane | 4 | Dunbar | Nebraska | – | – |
| **JJ-352** | *P. aceris* | 98.61 | Soybean | Rhizoplane | 6 | Dunbar | Nebraska | – | – |
| **JJ-355** | *P. illinoisensis* | 99.72 | Soybean | Rhizoplane | 6 | Dunbar | Nebraska | – | – |
| **JJ-364** | *P. urinalis* | 99.35 | Soybean | Rhizoplane | 7 | Dunbar | Nebraska | – | – |
| **JJ-365** | *P. odorifer* | 99.51 | Soybean | Rhizoplane | 7 | Dunbar | Nebraska | – | – |
| **JJ-372** | *P. aceris* | 99.30 | Soybean | Rhizoplane | 8 | Dunbar | Nebraska | – | – |
| **JJ-380** | *P. rigui* | 98.18 | Soybean | Rhizoplane | 10 | Dunbar | Nebraska | – | – |
| **JJ-385** | *P. silagei* | 97.09 | Soybean | Rhizoplane | 11 | Dunbar | Nebraska | – | – |
| **JJ-391** | *P. illinoisensis* | 99.65 | Soybean | Rhizoplane | 12 | Dunbar | Nebraska | – | – |
| **JJ-393** | *P. terrigena* | 99.17 | Soybean | Rhizoplane | 13 | Dunbar | Nebraska | – | – |
| **JJ-402** | *P. terrigena* | 99.79 | Soybean | Rhizoplane | 14 | Dunbar | Nebraska | – | – |
| **JJ-408** | *P. terrigena* | 99.04 | Soybean | Rhizoplane | 15 | Dunbar | Nebraska | – | – |
| **JJ-410** | *P. tritici* | 99.93 | Soybean | Rhizoplane | 15 | Dunbar | Nebraska | – | – |
| **JJ-411** | *P. aceris* | 98.68 | Soybean | Rhizoplane | 15 | Dunbar | Nebraska | – | – |
| **JJ-412** | *P. lautus* | 99.23 | Soybean | Rhizoplane | 15 | Dunbar | Nebraska | – | – |
| **JJ-413** | *P. 'taohuashanense'* | 98.80 | Soybean | Rhizoplane | 16 | Dunbar | Nebraska | – | – |
| **JJ-415** | *P. catalpae* | 99.59 | Soybean | Rhizoplane | 16 | Dunbar | Nebraska | – | – |
| **JJ-447** | *P. ehimensis* | 96.56 | Corn | Rhizoplane | 12 | Dunbar | Nebraska | – | – |
| **JJ-450** | *P. pectinilyticus* | 99.20 | Soybean | Rhizoplane | 5 | Dunbar | Nebraska | – | – |
| **JJ-460** | *P. pectinilyticus* | 99.78 | Corn | Endophyte | 9 | Dunbar | Nebraska | – | – |
| **JJ-467** | *P. pectinilyticus* | 98.85 | Soybean | Rhizoplane | 5 | Dunbar | Nebraska | – | – |
| **JJ-471** | *P. sabinae* | 99.46 | Soybean | Rhizoplane | 15 | Dunbar | Nebraska | – | – |
| **JJ-483** | *P. panacisoli* | 99.65 | Corn | Rhizoplane | 2 | Carrol | Iowa | – | – |
| **JJ-497** | *P. amylolyticus* | 99.59 | Corn | Rhizoplane | 3 | Carrol | Iowa | – | – |
| **JJ-514** | *P. dongdonensis* | 99.12 | Corn | Rhizoplane | 4 | Carrol | Iowa | – | – |
| **JJ-526** | *P. lautus* | 99.21 | Corn | Rhizoplane | 6 | Carrol | Iowa | – | – |

| | | | | | | | | | |
|---|---|---|---|---|---|---|---|---|---|
| **JJ-534** | *P. barcinonensis* | 99.11 | Corn | Rhizoplane | 7 | Carrol | Iowa | – | – |
| **JJ-539** | *P. cucumis* | 100.00 | Corn | Rhizoplane | 8 | Carrol | Iowa | – | – |
| **JJ-541** | *P. panacisoli* | 99.72 | Corn | Rhizoplane | 8 | Carrol | Iowa | – | – |
| **JJ-602** | *P. alvei* | 99.19 | Corn | Endophyte | 5 | Carrol | Iowa | – | – |
| **JJ-624** | *P. alkaliterrae* | 99.33 | Corn | Endophyte | 7 | Carrol | Iowa | – | – |
| **JJ-631** | *P. lautus* | 99.09 | Corn | Endophyte | 8 | Carrol | Iowa | – | – |
| **JJ-633** | *P. lautus* | 99.09 | Corn | Endophyte | 8 | Carrol | Iowa | – | – |
| **JJ-667** | *P. aceris* | 98.68 | Corn | Rhizoplane | 3 | Whitewater | Wisconsin | – | – |
| **JJ-676** | *P. 'taohuashanense'* | 98.59 | Corn | Rhizoplane | 5 | Whitewater | Wisconsin | – | – |
| **JJ-690** | *P. lautus* | 99.58 | Corn | Rhizoplane | 7 | Whitewater | Wisconsin | – | – |
| **JJ-702** | *P. tundrae* | 99.46 | Corn | Rhizoplane | 8 | Whitewater | Wisconsin | – | – |
| **JJ-718** | *P. 'taohuashanense'* | 98.73 | Corn | Rhizoplane | 10 | Whitewater | Wisconsin | – | – |
| **JJ-754** | *P. panacisoli* | 99.59 | Corn | Endophyte | 5 | Whitewater | Wisconsin | – | – |
| **JJ-764** | *P. castaneae* | 97.70 | Corn | Endophyte | 6 | Whitewater | Wisconsin | – | – |
| **JJ-773** | *P. amylolyticus* | 99.59 | Corn | Endophyte | 7 | Whitewater | Wisconsin | – | – |
| **JJ-778*** | *P. amylolyticus* | 99.80 | Corn | Endophyte | 8 | Whitewater | Wisconsin | – | – |
| **JJ-779** | *P. lactis* | 99.51 | Corn | Endophyte | 8 | Whitewater | Wisconsin | – | – |
| **JJ-782** | *P. amylolyticus* | 99.73 | Corn | Endophyte | 8 | Whitewater | Wisconsin | – | – |
| **JJ-788** | *P. amylolyticus* | 99.52 | Corn | Endophyte | 9 | Whitewater | Wisconsin | – | – |
| **JJ-794** | *P. lautus* | 99.44 | Corn | Endophyte | 10 | Whitewater | Wisconsin | – | – |
| **JJ-798** | *P. 'taohuashanense'* | 98.87 | Corn | Endophyte | 10 | Whitewater | Wisconsin | – | – |
| **JJ-816** | *P. barcinonensis* | 99.11 | Corn | Rhizoplane | 3 | Carrol | Iowa | – | – |
| **JJ-820** | *P. tundrae* | 99.59 | Corn | Rhizoplane | 6 | Carrol | Iowa | – | – |
| **JJ-824** | *P. illinoisensis* | 99.86 | Corn | Rhizoplane | 9 | Carrol | Iowa | – | – |
| **JJ-845** | *P. jamilae* | 99.52 | Corn | Endophyte | 10 | Whitewater | Wisconsin | + | – |
| **JJ-870** | *P. castaneae* | 99.32 | Soybean | Rhizoplane | 4 | Whitewater | Wisconsin | – | – |
| **JJ-890** | *P. contaminans* | 99.93 | Soybean | Rhizoplane | 7 | Whitewater | Wisconsin | – | – |
| **JJ-955** | *P. silagei* | 98.84 | Soybean | Endophyte | 7 | Whitewater | Wisconsin | – | – |
| **JJ-960** | *P. aestuarii* | 95.04 | Soybean | Endophyte | 8 | Whitewater | Wisconsin | – | – |
| **JJ-961** | *P. marinisediminis* | 99.52 | Soybean | Endophyte | 8 | Whitewater | Wisconsin | – | – |
| **JJ-963** | *P. tritici* | 99.57 | Soybean | Endophyte | 8 | Whitewater | Wisconsin | – | – |
| **JJ-964** | *P. lautus* | 99.44 | Soybean | Endophyte | 8 | Whitewater | Wisconsin | – | – |
| **JJ-985** | *P. castaneae* | 99.39 | Soybean | Rhizoplane | 2 | Whitewater | Wisconsin | – | – |
| **JJ-1000** | *P. xylanexedens* | 99.86 | Corn | Rhizoplane | 1 | Sparta | Illinois | – | – |
| **JJ-1004** | *P. barcinonensis* | 98.69 | Corn | Rhizoplane | 1 | Sparta | Illinois | – | – |
| **JJ-1057** | *P. amylolyticus* | 99.80 | Corn | Rhizoplane | 8 | Sparta | Illinois | – | – |
| **JJ-1059** | *P. lautus* | 99.23 | Corn | Rhizoplane | 8 | Sparta | Illinois | – | – |
| **JJ-1069** | *P. barcinonensis* | 98.69 | Corn | Endophyte | 1 | Sparta | Illinois | – | – |
| **JJ-1074** | *P. amylolyticus* | 99.86 | Corn | Endophyte | 1 | Sparta | Illinois | – | – |
| **JJ-1099** | *P. odorifer* | 98.58 | Corn | Endophyte | 6 | Sparta | Illinois | – | – |
| **JJ-1103** | *P. xylanilyticus* | 100.00 | Corn | Endophyte | 6 | Sparta | Illinois | – | – |
| **JJ-1115** | *P. amylolyticus* | 99.59 | Corn | Endophyte | 9 | Sparta | Illinois | – | – |
| **JJ-1181** | *P. terrigena* | 99.17 | Soybean | Rhizoplane | 8 | Sparta | Illinois | – | – |
| **JJ-1187** | *P. illinoisensis* | 99.79 | Soybean | Rhizoplane | 8 | Sparta | Illinois | – | – |
| **JJ-1277** | *P. dongdonensis* | 99.12 | Corn | Rhizoplane | 5 | Sparta | Illinois | – | – |
| **JJ-1283** | *P. pocheonensis* | 98.97 | Corn | Endophyte | 9 | Sparta | Illinois | – | – |

| | | | | | | | | | |
|---|---|---|---|---|---|---|---|---|---|
| **JJ-1311** | *P. amylolyticus* | 99.58 | Soybean | Endophyte | 6 | Sparta | Illinois | – | – |
| **JJ-1319** | *P. barcinonensis* | 99.17 | Soybean | Rhizoplane | 2 | Carrol | Iowa | – | – |
| **JJ-1343** | *P. barcinonensis* | 99.21 | Soybean | Rhizoplane | 5 | Carrol | Iowa | – | – |
| **JJ-1371** | *P. catalpae* | 99.66 | Soybean | Rhizoplane | 9 | Carrol | Iowa | – | – |
| **JJ-1402** | *P. susongensis* | 99.17 | Soybean | Endophyte | 2 | Carrol | Iowa | – | – |
| **JJ-1405** | *P. susongensis* | 99.17 | Soybean | Endophyte | 2 | Carrol | Iowa | – | – |
| **JJ-1410** | *P. odorifer* | 98.78 | Soybean | Endophyte | 3 | Carrol | Iowa | – | – |
| **JJ-1415** | *P. selenitireducens* | 99.23 | Soybean | Endophyte | 4 | Carrol | Iowa | – | – |
| **JJ-1425** | *P. granivorans* | 96.71 | Soybean | Endophyte | 5 | Carrol | Iowa | – | – |
| **JJ-1461** | *P. barcinonensis* | 99.24 | Corn | Rhizoplane | 1 | Troy | Ohio | – | – |
| **JJ-1465** | *P. cucumis* | 99.93 | Corn | Rhizoplane | 1 | Troy | Ohio | – | – |
| **JJ-1470** | *P. amylolyticus* | 99.45 | Corn | Rhizoplane | 2 | Troy | Ohio | – | – |
| **JJ-1476** | *P. massiliensis* | 99.86 | Corn | Rhizoplane | 2 | Troy | Ohio | – | – |
| **JJ-1487** | *P. pocheonensis* | 98.97 | Corn | Rhizoplane | 3 | Troy | Ohio | – | – |
| **JJ-1501** | *P. lautus* | 99.37 | Corn | Rhizoplane | 6 | Troy | Ohio | – | – |
| **JJ-1516** | *P. susongensis* | 99.24 | Corn | Rhizoplane | 8 | Troy | Ohio | – | – |
| **JJ-1522** | *P. barcinonensis* | 99.31 | Corn | Endophyte | 2 | Troy | Ohio | – | – |
| **JJ-1524** | *P. tritici* | 99.71 | Corn | Endophyte | 2 | Troy | Ohio | – | – |
| **JJ-1525** | *P. odorifer* | 99.93 | Corn | Endophyte | 2 | Troy | Ohio | – | – |
| **JJ-1531** | *P. rhizoplanae* | 98.21 | Corn | Endophyte | 3 | Troy | Ohio | – | – |
| **JJ-1544** | *P. barcinonensis* | 99.17 | Corn | Endophyte | 5 | Troy | Ohio | – | – |
| **JJ-1549** | *P. massiliensis* | 99.73 | Corn | Endophyte | 5 | Troy | Ohio | – | – |
| **JJ-1553** | *P. cucumis* | 99.79 | Corn | Endophyte | 6 | Troy | Ohio | – | – |
| **JJ-1555** | *P. barcinonensis* | 99.31 | Corn | Endophyte | 6 | Troy | Ohio | – | – |
| **JJ-1564** | *P. cucumis* | 99.59 | Corn | Endophyte | 7 | Troy | Ohio | – | – |
| **JJ-1570** | *P. susongensis* | 99.17 | Corn | Endophyte | 8 | Troy | Ohio | – | – |
| **JJ-1580** | *P. peoriae* | 99.72 | Soybean | Rhizoplane | 1 | Troy | Ohio | + | – |
| **JJ-1582** | *P. peoriae* | 99.65 | Soybean | Rhizoplane | 1 | Troy | Ohio | – | – |
| **JJ-1586** | *P. cucumis* | 100.00 | Soybean | Rhizoplane | 2 | Troy | Ohio | – | – |
| **JJ-1587** | *P. illinoisensis* | 99.72 | Soybean | Rhizoplane | 2 | Troy | Ohio | – | – |
| **JJ-1588** | *P. amylolyticus* | 99.52 | Soybean | Rhizoplane | 2 | Troy | Ohio | – | – |
| **JJ-1591** | *P. taichungensis* | 99.73 | Soybean | Rhizoplane | 2 | Troy | Ohio | – | – |
| **JJ-1595** | *P. odorifer* | 98.48 | Soybean | Rhizoplane | 3 | Troy | Ohio | – | – |
| **JJ-1596** | *P. barcinonensis* | 99.17 | Soybean | Rhizoplane | 3 | Troy | Ohio | – | – |
| **JJ-1598** | *P. odorifer* | 99.80 | Soybean | Rhizoplane | 3 | Troy | Ohio | – | – |
| **JJ-1601** | *P. illinoisensis* | 99.79 | Soybean | Rhizoplane | 3 | Troy | Ohio | – | – |
| **JJ-1602** | *P. typhae* | 98.18 | Soybean | Rhizoplane | 3 | Troy | Ohio | – | – |
| **JJ-1603** | *P. peoriae* | 99.86 | Soybean | Rhizoplane | 3 | Troy | Ohio | + | – |
| **JJ-1604** | *P. polymyxa* | 100.00 | Soybean | Rhizoplane | 3 | Troy | Ohio | + | – |
| **JJ-1611** | *P. typhae* | 97.08 | Soybean | Rhizoplane | 4 | Troy | Ohio | – | – |
| **JJ-1612** | *P. barcinonensis* | 99.24 | Soybean | Rhizoplane | 4 | Troy | Ohio | – | – |
| **JJ-1614** | *P. peoriae* | 99.65 | Soybean | Rhizoplane | 4 | Troy | Ohio | + | – |
| **JJ-1615** | *P. typhae* | 99.45 | Soybean | Rhizoplane | 4 | Troy | Ohio | – | – |
| **JJ-1620** | *P. massiliensis* | 99.86 | Soybean | Rhizoplane | 5 | Troy | Ohio | – | – |
| **JJ-1623** | *P. dongdonensis* | 99.12 | Soybean | Rhizoplane | 5 | Troy | Ohio | – | – |
| **JJ-1631** | *P. dongdonensis* | 99.27 | Soybean | Rhizoplane | 6 | Troy | Ohio | – | – |
| **JJ-1638** | *P. peoriae* | 99.72 | Soybean | Rhizoplane | 7 | Troy | Ohio | + | + |

| | | | | | | | | | |
|---|---|---|---|---|---|---|---|---|---|
| **JJ-1639** | *P. cucumis* | 100.00 | Soybean | Rhizoplane | 7 | Troy | Ohio | – | – |
| **JJ-1640** | *P. peoriae* | 99.72 | Soybean | Rhizoplane | 7 | Troy | Ohio | + | – |
| **JJ-1648** | *P. turicensis* | 97.36 | Soybean | Rhizoplane | 8 | Troy | Ohio | – | – |
| **JJ-1650** | *P. jamilae* | 99.88 | Soybean | Rhizoplane | 8 | Troy | Ohio | + | – |
| **JJ-1652** | *P. peoriae* | 99.72 | Soybean | Rhizoplane | 9 | Troy | Ohio | + | – |
| **JJ-1653** | *P. cucumis* | 100.00 | Soybean | Rhizoplane | 9 | Troy | Ohio | – | – |
| **JJ-1667** | *P. odorifer* | 99.86 | Soybean | Endophyte | 1 | Troy | Ohio | – | – |
| **JJ-1669** | *P. odorifer* | 98.58 | Soybean | Endophyte | 1 | Troy | Ohio | – | – |
| **JJ-1678** | *P. cucumis* | 100.00 | Soybean | Endophyte | 2 | Troy | Ohio | – | – |
| **JJ-1679** | *P. cucumis* | 100.00 | Soybean | Endophyte | 2 | Troy | Ohio | – | – |
| **JJ-1680** | *P. massiliensis* | 99.86 | Soybean | Endophyte | 2 | Troy | Ohio | – | – |
| **JJ-1683*** | *P. jamilae* | 99.73 | Soybean | Endophyte | 3 | Troy | Ohio | + | – |
| **JJ-1684** | *P. cucumis* | 99.31 | Soybean | Endophyte | 3 | Troy | Ohio | – | – |
| **JJ-1688** | *P. massiliensis* | 99.86 | Soybean | Endophyte | 3 | Troy | Ohio | – | – |
| **JJ-1692** | *P. typhae* | 99.59 | Soybean | Endophyte | 4 | Troy | Ohio | – | – |
| **JJ-1693** | *P. amylolyticus* | 99.52 | Soybean | Endophyte | 4 | Troy | Ohio | – | – |
| **JJ-1701** | *P. typhae* | 99.66 | Soybean | Endophyte | 5 | Troy | Ohio | – | – |
| **JJ-1703** | *P. typhae* | 99.59 | Soybean | Endophyte | 5 | Troy | Ohio | – | – |
| **JJ-1709** | *P. massiliensis* | 99.93 | Soybean | Endophyte | 5 | Troy | Ohio | – | – |
| **JJ-1715** | *P. peoriae* | 99.72 | Soybean | Endophyte | 6 | Troy | Ohio | + | + |
| **JJ-1720** | *P. peoriae* | 99.72 | Soybean | Endophyte | 6 | Troy | Ohio | + | + |
| **JJ-1722*** | *P. peoriae* | 99.72 | Soybean | Endophyte | 7 | Troy | Ohio | + | + |
| **JJ-1723** | *P. dongdonensis* | 99.27 | Soybean | Endophyte | 7 | Troy | Ohio | – | – |
| **JJ-1724** | *P. peoriae* | 99.72 | Soybean | Endophyte | 7 | Troy | Ohio | + | + |
| **JJ-1725** | *P. amylolyticus* | 99.66 | Soybean | Endophyte | 7 | Troy | Ohio | – | – |
| **JJ-1729** | *P. polymyxa* | 100.00 | Soybean | Endophyte | 8 | Troy | Ohio | – | – |
| **JJ-1736** | *P. typhae* | 99.59 | Soybean | Endophyte | 8 | Troy | Ohio | – | – |
| **JJ-1742** | *P. cucumis* | 100.00 | Soybean | Endophyte | 9 | Troy | Ohio | – | – |
| **JJ-1743** | *P. polymyxa* | 99.93 | Soybean | Endophyte | 9 | Troy | Ohio | + | – |
| **JJ-1744** | *P. typhae* | 99.65 | Soybean | Endophyte | 9 | Troy | Ohio | – | – |
| **JJ-1747** | *P. polymyxa* | 99.65 | Soybean | Endophyte | 10 | Troy | Ohio | + | + |
| **JJ-1755** | *P. graminis* | 100.00 | Soybean | Rhizoplane | 5 | Carrol | Iowa | – | – |
| **JJ-1759** | *P. catalpae* | 99.66 | Soybean | Rhizoplane | 8 | Carrol | Iowa | – | – |
| **JJ-1762** | *P. alginolyticus* | 99.29 | Soybean | Endophyte | 4 | Carrol | Iowa | – | – |
| **JJ-1774** | *P. susongensis* | 99.21 | Corn | Rhizoplane | 2 | Troy | Ohio | – | – |
| **JJ-1781** | *P. barcinonensis* | 99.03 | Corn | Rhizoplane | 5 | Troy | Ohio | – | – |
| **JJ-1783** | *P. sonchi* | 98.64 | Corn | Rhizoplane | 6 | Troy | Ohio | – | – |
| **JJ-1789** | *P. dongdonensis* | 99.27 | Soybean | Rhizoplane | 2 | Troy | Ohio | – | – |
| **JJ-1817** | *P. rhizoplanae* | 98.35 | Corn | Endophyte | 6 | Troy | Ohio | – | – |
| **JJ-1819** | *P. contaminans* | 99.93 | Soybean | Endophyte | 1 | Troy | Ohio | – | – |
| **JJ-1831** | *P. typhae* | 97.33 | Soybean | Endophyte | 9 | Troy | Ohio | – | – |
| **JM-929** | *P. odorifer* | 99.51 | Cotton | Endophyte | n.a | Tallassee | Alabama | – | – |
| **JM-1337** | *P. cucumis* | 99.86 | n.a | Endophyte | n.a | Tallassee | Alabama | – | – |
| **JM-1355** | *P. barcinonensis* | 99.16 | n.a | Endophyte | n.a | Tallassee | Alabama | – | – |
| **JM-1368** | *P. barcinonensis* | 99.07 | n.a | Endophyte | n.a | Tallassee | Alabama | – | – |
| **JM-1419** | *P. cucumis* | 99.86 | n.a | Endophyte | n.a | Tallassee | Alabama | – | – |
| **JM-1420** | *P. cucumis* | 99.86 | n.a | Endophyte | n.a | Tallassee | Alabama | – | – |

| | | | | | | | | | |
|---|---|---|---|---|---|---|---|---|---|
| **AP-66** | *P. silvae* | 99.50 | n.a | n.a | n.a | n.a | n.a | – | – |

* Whole genome sequence is available

**Table S2. List of *Paenibacillus* type strains used for the phylogenetic tree construction.**

| Strain | GenBank accession number |
|---|---|
| *P. massiliensis subsp. panacisoli* DSM 21345 | AB245384.1 |
| *P. ehimensis* DSM 11029 | AY116665.1 |
| *P. illinoisensis* DSM 11733 | NR_115624.1 |
| *P. selenitireducens* KCTC 33157 | NR_133807.1 |
| *P. cucumis* DSM 101601 | NR_149778.1 |
| *P. thiaminolyticus* DSM 7262 | AB073197.1 |
| *P. terrigena* DSM 21567 | AB248087.1 |
| *P. granivorans* | AF237682.1 |
| *P. turicensis* DSM 14349 | AF378694.1 |
| *P. agaridevorans* DSM 1355 | AJ345023.1 |
| *P. cineris* DSM 16945 | AJ575658.1 |
| *P. barcinonensis* DSM 15478 | AJ716019.1 |
| *P. lactis* DSM 15596 | AY257868.1 |
| *P. xylanilyticus* DSM 17255 | AY427832.1 |
| *P. alkaliterrae* DSM 17040 | AY960748.1 |
| *P. urinalis* DSM 22281 | EF212892.1 |
| *P. contaminans* BCRC 17728 | EF626690.1 |
| *P. taichungensis* DSM 19942 | EU179327.1 |
| *P. peoriae* DSM 8320 | EU391157.1 |
| *P. xylanexedens* DSM 21292 | EU558281.1 |
| *P. tundrae* DSM 21291 | EU558284.1 |
| *P. typhae* DSM 25190 | NR_109462.1 |
| *P. pocheonensis* DSM 23906 | NR_112565.1 |
| *P. alginolyticus DSM 5050* | NR_115595.1 |
| *P. sonchi* CCBAU 83901 | NR_115751.1 |
| *P. aestuarii* DSM 23861 | NR_116365.1 |
| *P. catalpae* DSM 24714 | NR_118012.1 |
| *P. taohuashanense* DSM 25809 | NR_118393.1 |
| *P. sabinae* T27 | NR_121732.2 |
| *P. dongdonensis* DSM 27607 | NR_134112.1 |
| *P. susongensis* JCM 19951 | NR_134118.1 |
| *P. endophyticus* LMG 27297 | NR_135705.1 |
| *P. oenotherae* JCM 19573 | NR_136822.1 |
| *P. etheri* DSM 29760 | NR_148622.1 |
| *P. aceris* DSM 24950 | NR_156841.1 |
| *P. rhizoplanae* DSM 103963 | NR_156842.1 |
| *P. tritici* CECT 9125 | NR_157638.1 |
| *P. solisilvae* JCM 32513 | NR_175458.1 |
| *P. lautus* DSM 3035 | NZ_BIMF01000051.1 |
| *P. amylolyticus* NBRC 15957 | NZ_BIMJ01000009.1 |
| *P. rigui* WPCB173 | NZ_NMQW01000006.1 |

**Table S3. Annotation of the known specialized metabolites detected in the extracts of 227 plant-associated *Paenibacillus* isolates. For the metabolites highlighted in bold, annotation was performed on the MS/MS level.**

| Parent ion *m/z* | Adduct ion | Compound | Family | Bioactivity |
|---|---|---|---|---|
| 442.2839 | $[M+2H]^{2+}$ | Fusaricidin A | Fusaricidins | Gram +, Fungi (Kajimura & Kaneda, 1996) |
| **449.2909** | $[M+2H]^{2+}$ | Fusaricidin B/Antibiotic LI-F05A/LI-F06A | Fusaricidins | Gram +, Fungi (Ryu *et al.*, 2017) |
| 449.2921 | $[M+2H]^{2+}$ | Fusaricidin B/Antibiotic LI-F05A/LI-F06A | Fusaricidins | Gram +, Fungi (Ryu *et al.*, 2017) |
| 456.300 | $[M+2H]^{2+}$ | Antibiotic LI-F05B/LI-F06B/LI-F08A | Fusaricidins | Gram +, Fungi (Ryu *et al.*, 2017) |
| 463.3076 | $[M+2H]^{2+}$ | Antibiotic LI-F08B | Fusaricidins | Gram +, Fungi (Ryu *et al.*, 2017) |
| 463.3077 | $[M+2H]^{2+}$ | Antibiotic LI-F08B | Fusaricidins | Gram +, Fungi (Ryu *et al.*, 2017) |
| 466.2835 | $[M+2H]^{2+}$ | Antibiotic LI-F07A | Fusaricidins | Gram +, Fungi (Kajimura & Kaneda, 1996) |
| 466.2848 | $[M+2H]^{2+}$ | Antibiotic LI-F07A | Fusaricidins | Gram +, Fungi (Kajimura & Kaneda, 1996) |
| 473.2915 | $[M+2H]^{2+}$ | Antibiotic LI-F07B | Fusaricidins | Gram +, Fungi (Ryu *et al.*, 2017) |
| 481.2895 | $[M+2H]^{2+}$ | Fusaricidine D | Fusaricidins | Gram +, Fungi (Ryu *et al.*, 2017) |
| 883.5606 | $[M+H]^{+}$ | Fusaricidin A | Fusaricidins | Gram +, Fungi (Kajimura & Kaneda, 1996) |
| 911.5916 | $[M+H]^{+}$ | Antibiotic LI-F08A | Fusaricidins | Gram +, Fungi (Kajimura & Kaneda, 1996) |
| 911.5921 | $[M+H]^{+}$ | Antibiotic LI-F08A | Fusaricidins | Gram +, Fungi (Kajimura & Kaneda, 1996) |
| 925.6076 | $[M+H]^{+}$ | Antibiotic LI-F08B | Fusaricidins | Gram +, Fungi (Ryu *et al.*, 2017) |
| 945.5767 | $[M+H]^{+}$ | Antibiotic LI-F07B | Fusaricidins | Gram +, Fungi (Ryu *et al.*, 2017) |
| 947.5557 | $[M+H]^{+}$ | Fusaricidin C | Fusaricidins | Gram +, Fungi |

| | | | | |
|---|---|---|---|---|
| | | | | (Kajimura & Kaneda, 1996) |
| 961.5712 | $[M+H]^+$ | Fusaricidine D | Fusaricidins | Gram +, Fungi (Ryu *et al.*, 2017) |
| 961.5718 | $[M+H]^+$ | Fusaricidine D | Fusaricidins | Gram +, Fungi (Ryu *et al.*, 2017) |
| 381.9119 | $[M+3H]^{3+}$ | Polymyxin M2/A1 | Polymyxins | Gram – (Martin *et al.*, 2003) |
| 385.9241 | $[M+3H]^{3+}$ | Polymyxin E | Polymyxins | Gram – (Ikai *et al.*, 1998) |
| 386.5838 | $[M+3H]^{3+}$ | Polymyxin M1/A2 | Polymyxins | Gram – (Martin *et al.*, 2003) |
| 386.5839 | $[M+3H]^{3+}$ | Polymyxin M1/A2 | Polymyxins | Gram – (Martin *et al.*, 2003) |
| 390.5958 | $[M+3H]^{3+}$ | Polymyxin E | Polymyxins | Gram – (Ikai *et al.*, 1998) |
| **390.596** | $[M+3H]^{3+}$ | Polymyxin E | Polymyxins | Gram – (Ikai *et al.*, 1998) |
| 397.9119 | $[M+3H]^{3+}$ | Polymyxin P1 | Polymyxins | Gram – (Niu *et al.*, 2013) |
| 401.9238 | $[M+3H]^{3+}$ | Polymyxin B | Polymyxins | Gram – (Pittenauer *et al.*, 2006) |
| 579.371 | $[M+2H]^{2+}$ | Polymyxin M1 | Polymyxins | Gram – (Martin *et al.*, 2003) |
| 579.3713 | $[M+2H]^{2+}$ | Polymyxin M1 | Polymyxins | Gram – (Martin *et al.*, 2003) |
| 585.3898 | $[M+2H]^{2+}$ | Polymyxin E7 | Polymyxins | Gram – (Ikai *et al.*, 1998) |
| 596.3656 | $[M+2H]^{2+}$ | Polymyxin P1 | Polymyxins | Gram – (Niu *et al.*, 2013) |
| 602.382 | $[M+2H]^{2+}$ | Polymyxin B5 | Polymyxins | Gram – (Pittenauer *et al.*, 2006) |
| 610.3812 | $[M+2H]^{2+}$ | Polymyxin B6 | Polymyxins | Gram – (Pittenauer *et al.*, 2006) |
| 486.9491 | $[M+3H]^{3+}$ | Tridecaptin $B_1$ | Tridecaptins | Gram – (Cochrane *et al.*, 2015) |
| **496.9524** | $[M+3H]^{3+}$ | Tridecaptin M | Tridecaptins | Gram – |

| | | | | |
|---|---|---|---|---|
| | | | | (Jangra *et al.*, 2019b) |
| 744.9239 | $[M+2H]^{2+}$ | Tridecaptin M | Tridecaptins | Gram – (Jangra *et al.*, 2019b) |
| 789.9477 | $[M+2H]^{2+}$ | Tridecaptin $A_3$ | Tridecaptins | Gram – (Lohans *et al.*, 2012) |
| 539.9715 | $[M+3H]^{3+}$ | Tridecaptin $A_5$ | Tridecaptins | Gram -, Gram + |

**Table S4. NMR data of paenitracin A (1), measured in DMSO-*d*$_6$ at 298 K*.**

| | Residue | NH | $H_\alpha$($C_\alpha$, type) | $H_\beta$($C_\beta$, type) | Other |
|---|---|---|---|---|---|
| **1** | **1-Ile** | ND | 3.55,d (57.0, CH) | 1.66 (39.5, CH) | 1 C: ND<br>4 $CH_2$: 1.41, 1.15 (25.5)<br>5 $CH_3$: 0.84 (11.3)<br>6 $CH_3$: 0.81 (13.3) |
| **2** | **2-Cys** | | 5.13 (78.5, CH) | 3.41 (32.7, $CH_2$) | |
| **3** | **3-Leu** | 7.75 | 4.35 (51.9, CH) | 1.52, 1.46 (41.0, $CH_2$) | 4 CH: 1.54 (24.0)<br>5 $CH_3$: 0.85 (21.2)<br>6 $CH_3$: 0.89 (22.8) |
| **4** | **4-Glu** | 8.18 | 4.06 (53.6, CH) | 1.81, 1.75 (27.7, $CH_2$) | 4 $CH_2$: 2.19 (32.8, $CH_2$) |
| **5** | **5-Ile** | 7.22 | 3.95 (57.6, CH) | 1.75 (35.4, CH) | 4 $CH_2$: 1.46, 1.17 (24.3)<br>5 $CH_3$: 0.78 (10.8)<br>6 $CH_3$: 0.94 (15.0) |
| **6** | **6-Lys** | 8.46 | 4.49 (52.9, CH) | 1.42, 1.15 (25.5, $CH_2$) | 4 $CH_2$: 1.54 (24.0)<br>5 $CH_2$: 1.47 (24.3)<br>6 $CH_2$: 2.95, 2.82 (36.8)<br>NH: 7.52 |
| **7** | **7-Leu** | 10.37 | 4.22 (49.8, CH) | 1.61, 1.36 (38.7, $CH_2$) | 4 CH: 1.51 (24.1)<br>5 $CH_3$: 0.77 (20.4)<br>6 $CH_3$: 0.85 (22.9) |
| **8** | **8-Ile** | 7.09 | 4.26 (55.6, CH) | 1.68 (37.5, CH) | 4 $CH_2$: 1.28, 0.70 (23.9)<br>5 $CH_3$: 0.70 (11.5)<br>6 $CH_3$: 0.66 (14.1) |
| **9** | **9-Trp** | 8.98 | 5.18 (52.2, CH) | 3.11, 2.83 (28.5, $CH_2$) | 4 C: ND<br>5 CH: 7.18 (123.2)<br>6 NH: 10.9<br>7 C: 135.5<br>8 CH: 7.26 (110.7)<br>9 CH: 7.01 (120.3)<br>10 CH: 6.95 (117.7)<br>11 CH: 7.48 (117.8)<br>12 C: 126.7 |
| **10** | **10-Thr** | 10.78 | 3.66 (60.4, CH) | 4.02 (66.8, CH) | 4 $CH_3$: 0.67 (18.5) |
| **11** | **11-Asp** | 8.84 | 4.09 (50.2, CH) | 2.43, 2.01 (38.2, $CH_2$) | |
| **12** | **12-Asn** | 7.82 | 3.94 (51.8, CH) | 2.38, 2.18 (35.2, $CH_2$) | $NH_2$: 7.58, 6.74 |

* $^{1}$H 850 MHz and $^{13}$C chemical shifts inferred from HSQC and HMBC spectra

ND: not determined under these experimental conditions (and also all carbonyl carbons).

**Table S5.** Retention times ($t_R$, min) of the L-FDAA derivatives for natural paenitracin A (**1**) and standard amino acids.

| | **[M+H]⁺** | **$t_R$, min** | | | |
|---|---|---|---|---|---|
| | | L-AA (standard) | D-AA (standard) | Paen A | Stereochemical assignment |
| **Asp** | 386.0943 | 4.75 | 4.86 | 4.75/4.86 | **L + D** |
| **Cys*** | 626.126 | 6.37 | 6.62 | - | **NA** |
| **Glu** | 400.1099 | 4.92 | 5.07 | 5.07 | **D** |
| **Ile** | 384.1514 | 6.40 | 6.93 | 6.40 | **L** |
| **Leu** | 384.1514 | 6.50 | 7.01 | 6.50/7.01 | **L + D** |
| **Lys*** | 651.2118 | 6.45 | 6.63 | 6.45 | **L** |
| **Thr** | 372.115 | 4.75 | 5.12 | 4.75 | **L** |
| **Trp** | 457.1466 | 6.31 | 6.58 | - | **NA** |

* Product of double-addition of Marfey`s reagent

NA: not available

**Table S6.** Retention times ($t_R$, min) of the L-FDAA derivatives for natural paenitracin B (**2**) and standard amino acids.

| | **[M+H]⁺** | **$t_R$, min** | | | |
|---|---|---|---|---|---|
| | | L-AA (standard) | D-AA (standard) | Paen B | Stereochemical assignment |
| **Asp** | 386.0943 | 4.75 | 4.86 | 4.75/4.86 | **L + D** |
| **Cys*** | 626.126 | 6.37 | 6.62 | - | **NA** |
| **Glu** | 400.1099 | 4.92 | 5.07 | 5.07 | **D** |
| **Ile** | 384.1514 | 6.40 | 6.93 | 6.40 | **L** |
| **Leu** | 384.1514 | 6.50 | 7.01 | 6.50/7.01 | **L + D** |
| **Lys*** | 651.2118 | 6.45 | 6.63 | 6.45 | **L** |
| **Thr** | 372.115 | 4.75 | 5.12 | 4.75 | **L** |
| **Trp** | 457.1466 | 6.31 | 6.58 | - | **NA** |

* Product of double-addition of Marfey`s reagent

NA: not available

**Table S7.** Antagonization of in vitro antibacterial activity of paenitracin A against *S. aureus* ATCC 29213 by addition of $C_{10}$PP.

| **Antagonist** | **$ZnSO_4$** | ***S. aureus* ATCC 29213** |
|---|---|---|
| **$C_{10}$PP** | 0.3 mM | G |
| **-** | 0.3 mM | NG |
| **$C_{10}$PP** | - | G |
| **-** | - | G |

Experiments were performed in triplicates. NG = no visible bacterial growth, corresponding to unaffected antibiotic activity. G = visible bacterial growth, corresponding to antagonization of antibiotic activity.

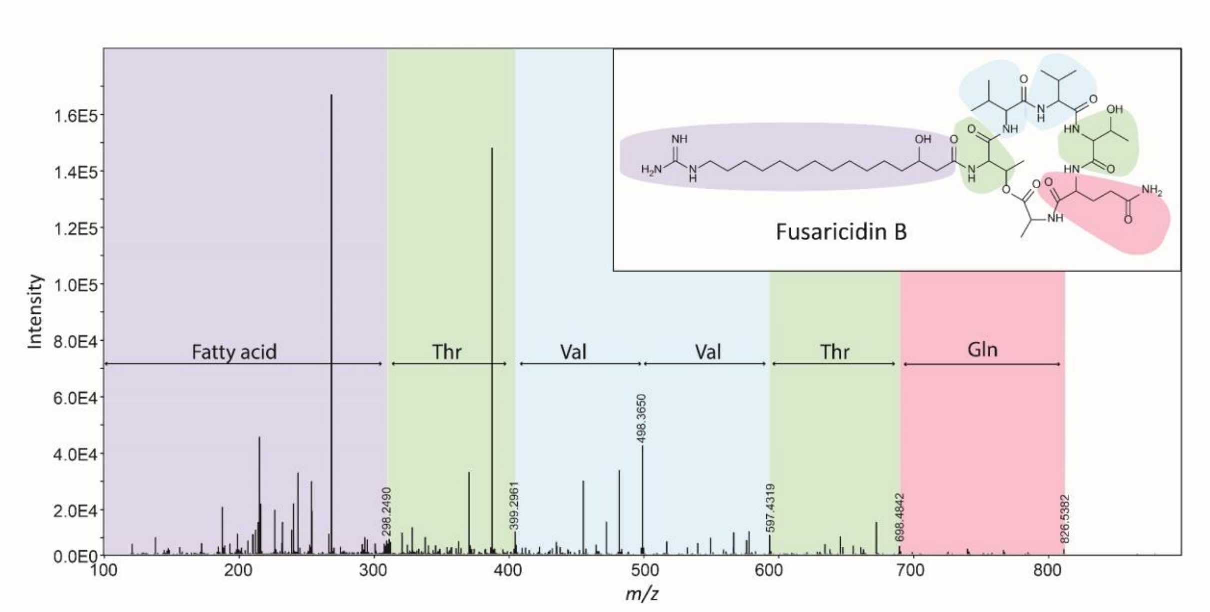


**Figure S1.** MS/MS spectrum of fusaricidin B (precursor ion $[M + 2H]^{2+}$ *m/z* 449.2909). The assignment of the sequence of amino acid residues is based on the mass differences between the consecutive *b* ions.

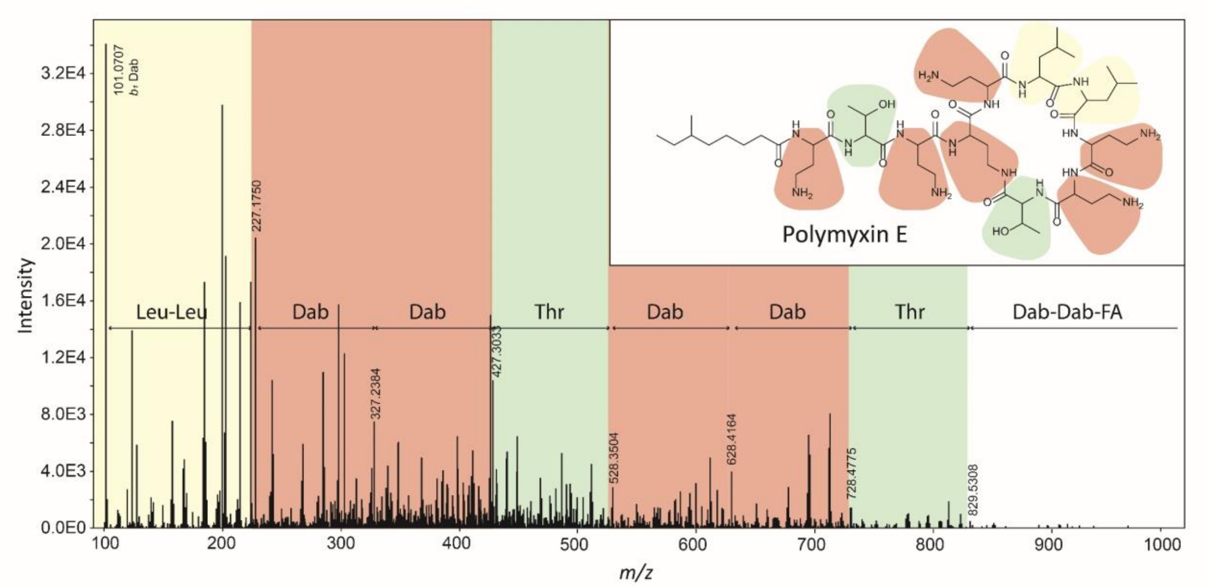


**Figure S2.** MS/MS spectrum of polymyxin E (precursor ion $[M + 3H]^{3+}$ *m/z* 390.596). The assignment of the sequence of amino acid residues is based on the mass differences between the consecutive *b* ions.

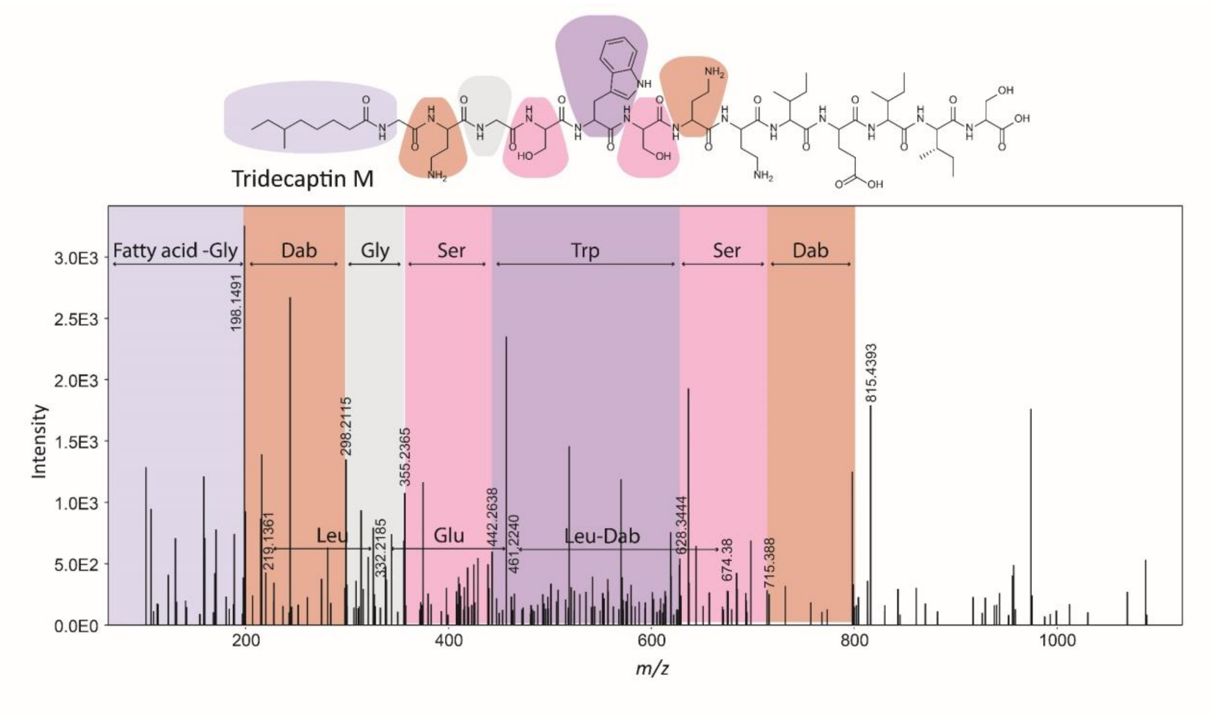


**Figure S3.** MS/MS spectrum of tridecaptin M (precursor ion $[M + 3H]^{3+}$ *m/z* 496.9524). The assignment of the sequence of amino acid residues is based on the mass differences between the consecutive *b* ions.

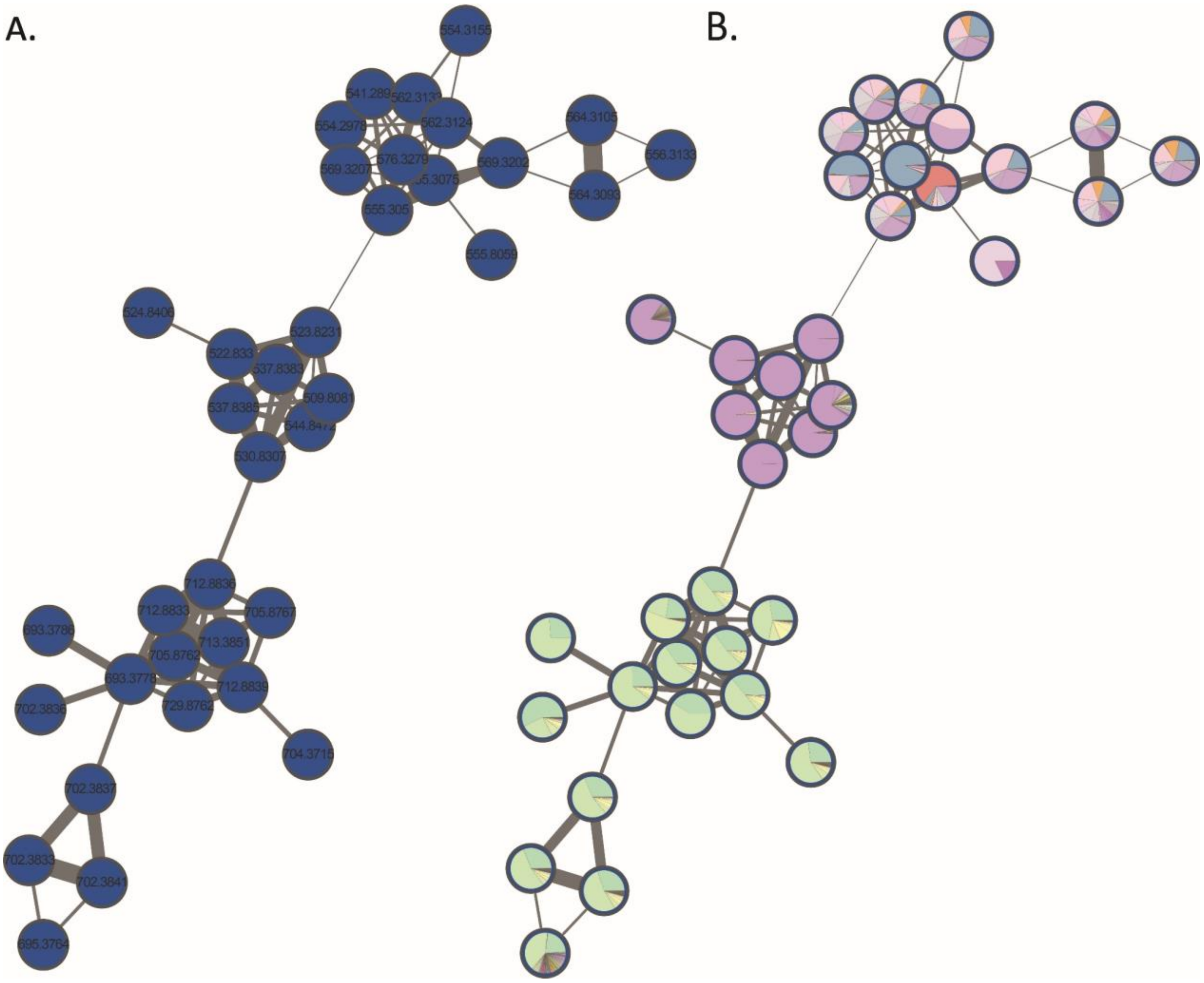


**Figure S4. A.** The MassQL-annotated molecular family of lysine-containing natural products. **B**. On the same molecular family, pie charts were mapped to the nodes to represent the relative precursor ion intensities within each extract.

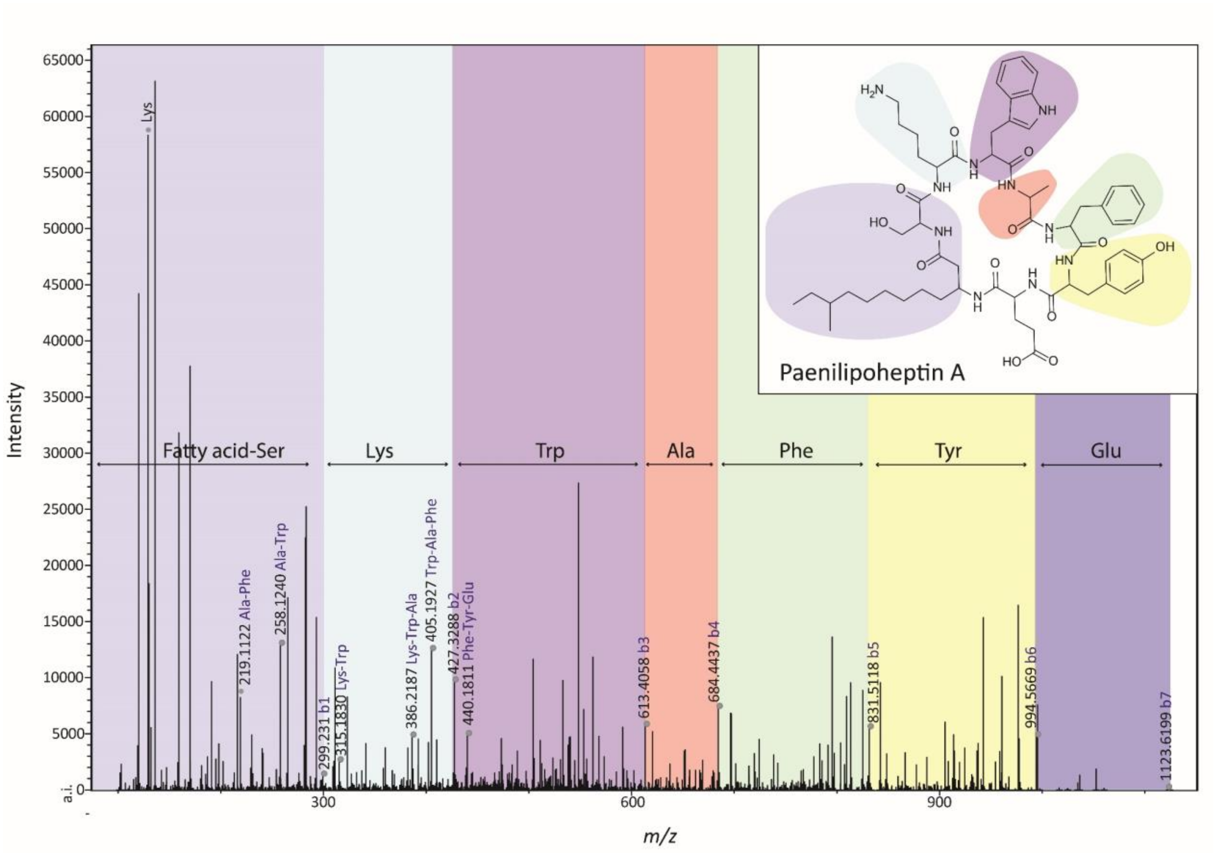


**Figure S5.** MS/MS spectrum of paenilipoheptin A (precursor ion $[M+2H]^{2+}$ *m/z* 562.3130). The assignment of the sequence of amino acid residues is based on the mass differences between the consecutive *b* ions.

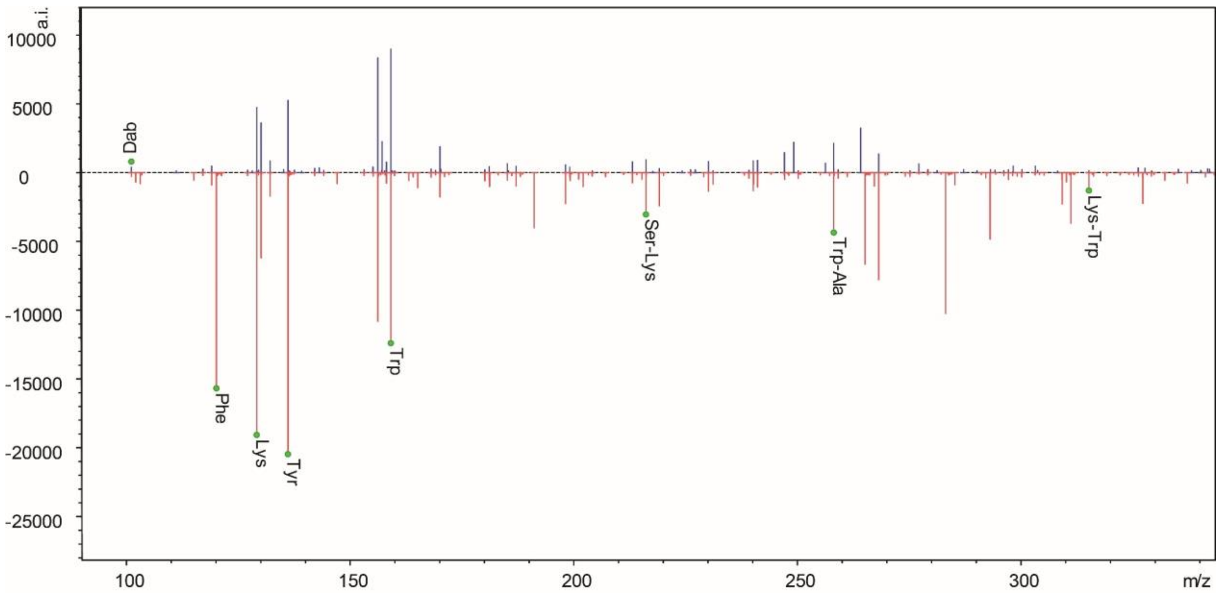


**Figure S6.** Direct MS/MS spectra comparison of the mass feature having *m/z* of 555.305 (red) with the mass feature having *m/z* of 523.8231 (blue).

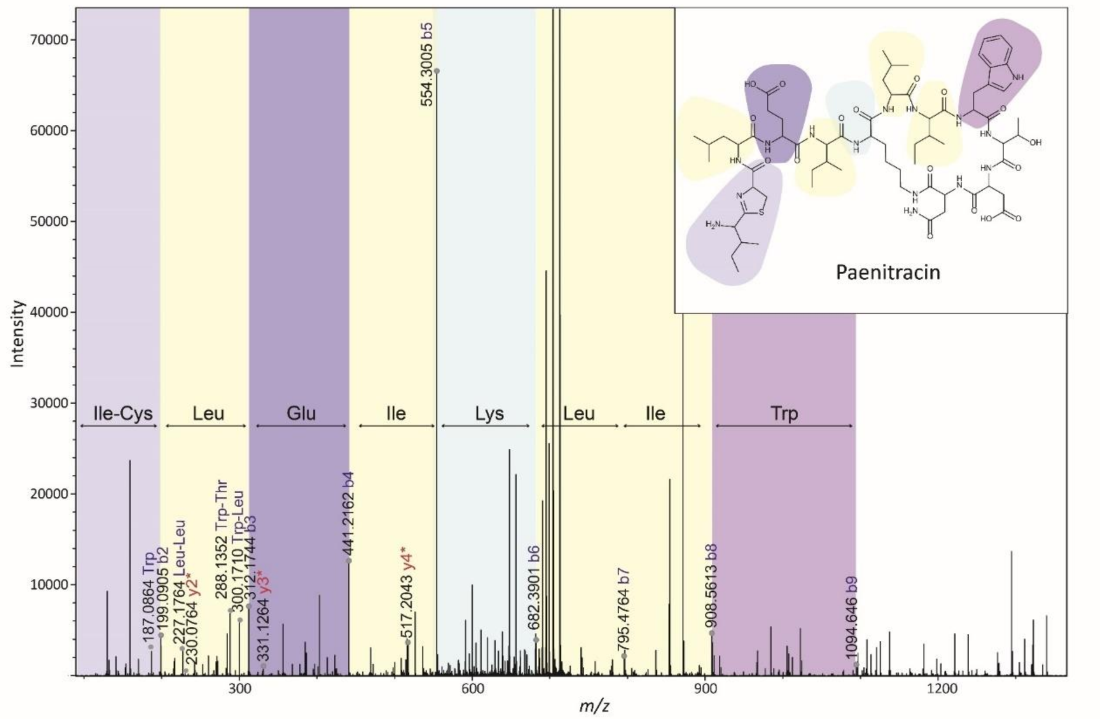


**Figure S7.** MS/MS spectrum of paenitracin A (precursor ion $[M + 2H]^{2+}$ *m/z* 712.8843). The assignment of the sequence of amino acid residues is based on the mass differences between the consecutive *b* ions.

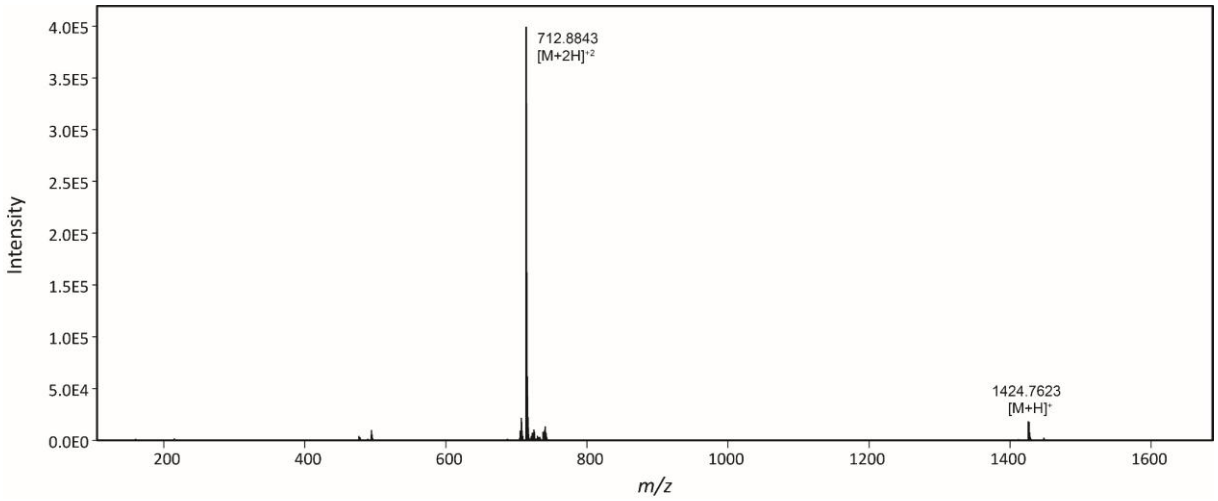


**Figure S8**. HRMS spectrum of **1**.

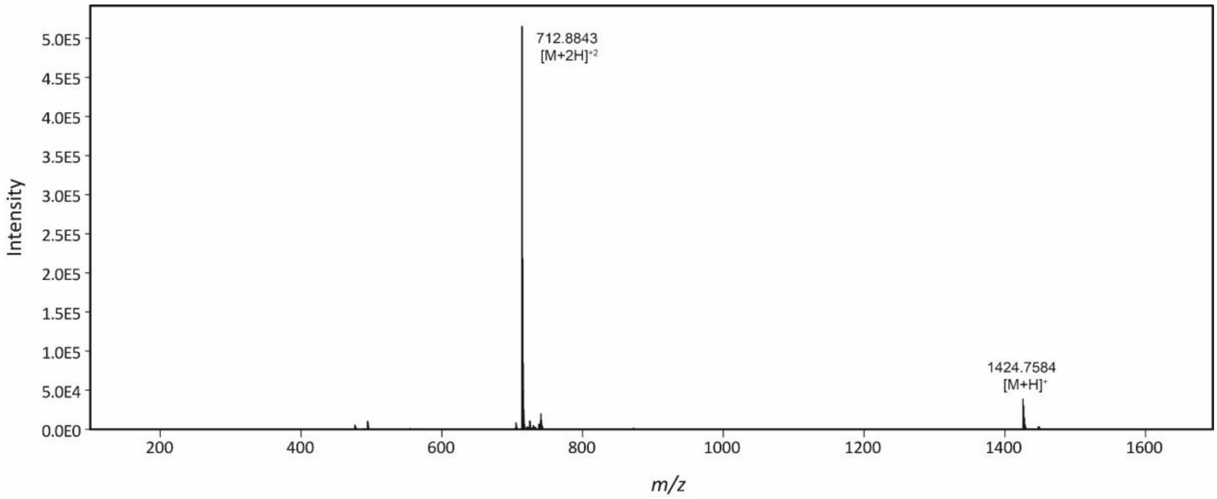


**Figure S9**. HRMS spectrum of **2**.

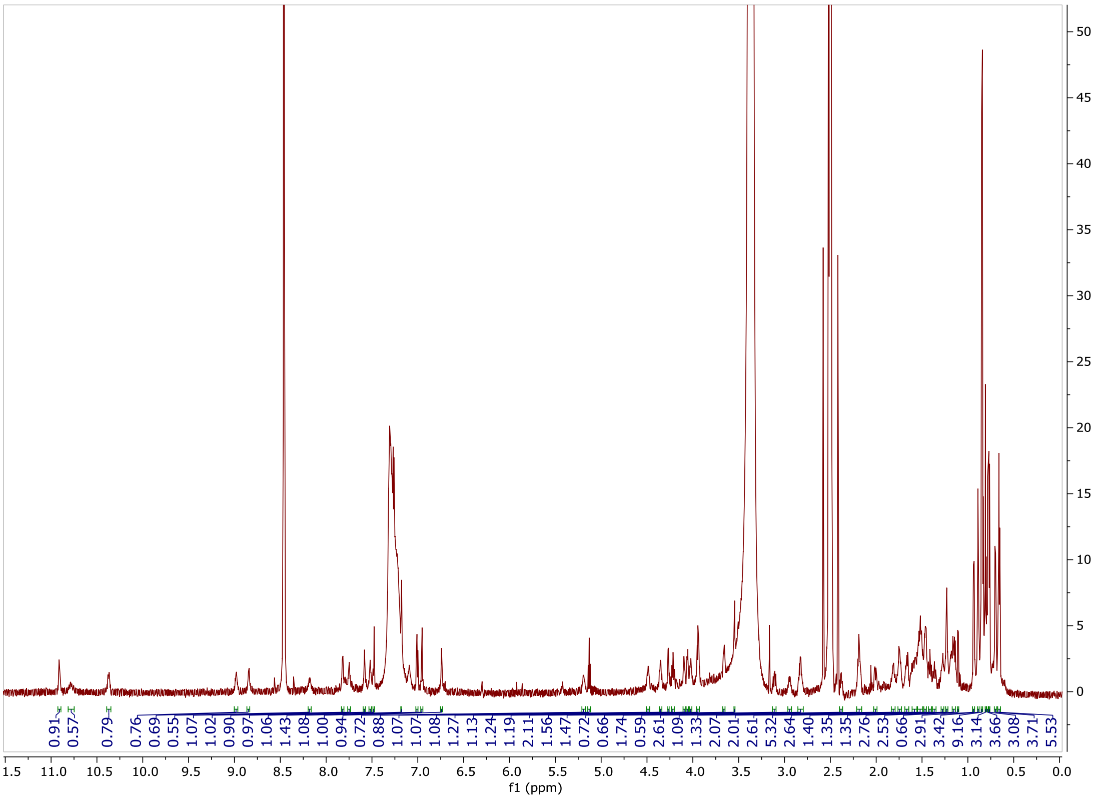


**Figure S10**. $^{1}H$ NMR spectrum of **1** (850 MHz, in DMSO-$d_6$).

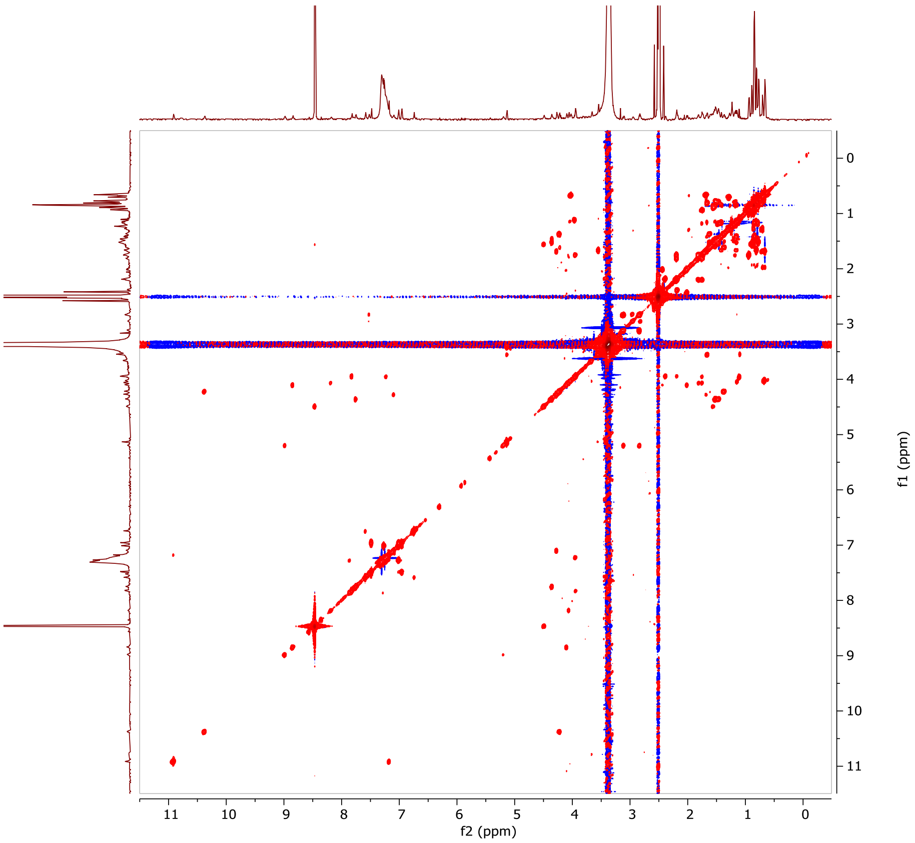


**Figure S11**. $^{1}H$–$^{1}H$ COSY spectrum of **1** (850 MHz, in DMSO-$d_6$).

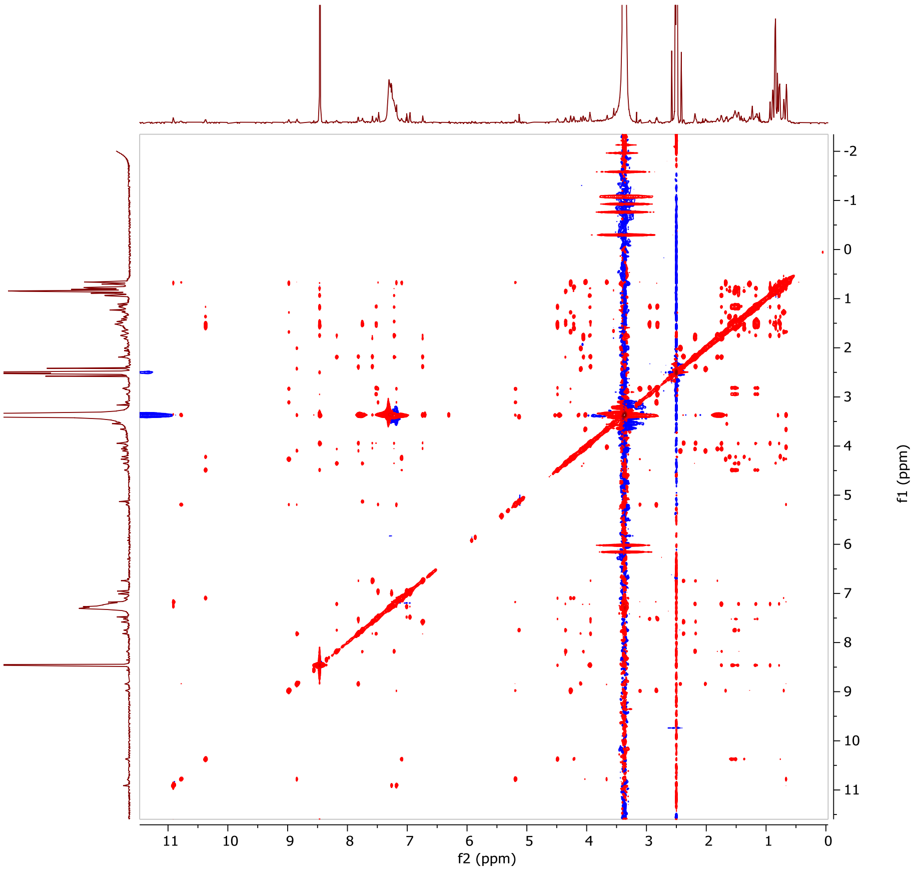


**Figure S12**. NOESY spectrum of **1** (850 MHz, in DMSO-$d_6$).

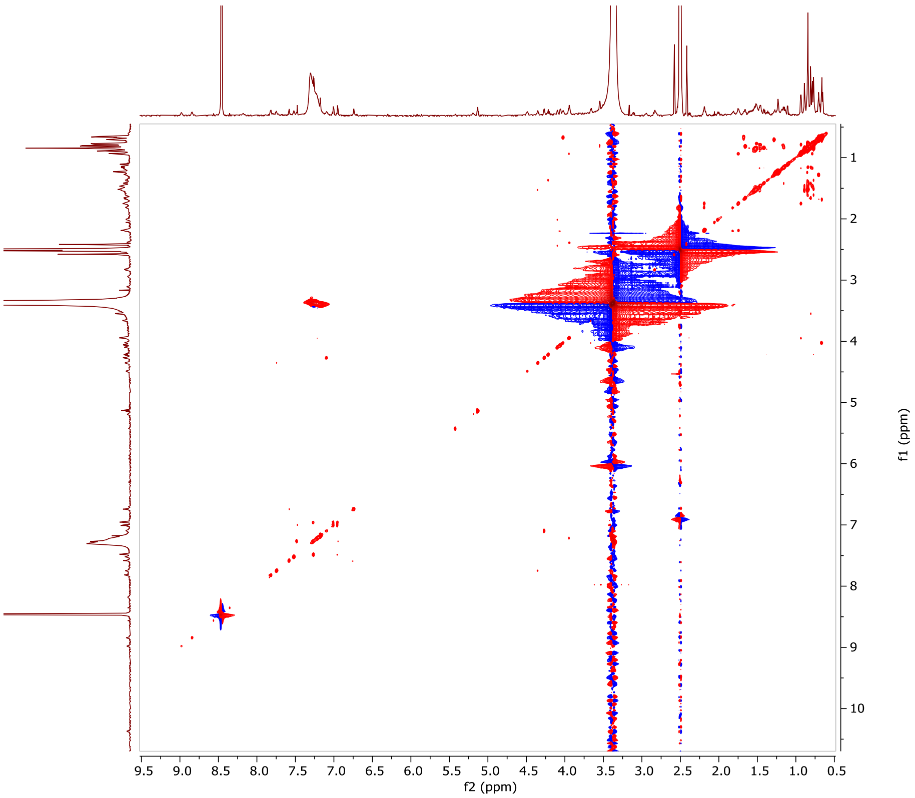


**Figure S13**. $^{1}H$–$^{1}H$ TOCSY spectrum of **1** (850 MHz, in DMSO-$d_6$).

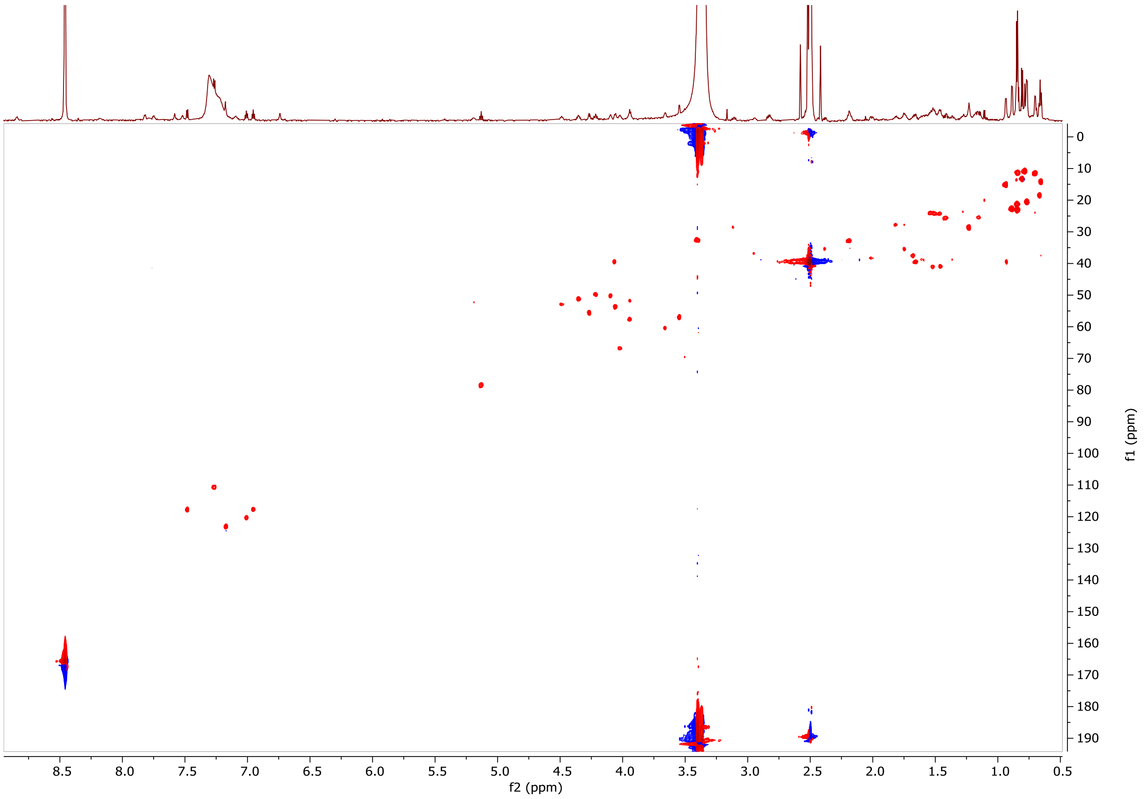


**Figure S14**. HSQC spectrum of **1** (850 MHz, in DMSO-$d_6$).

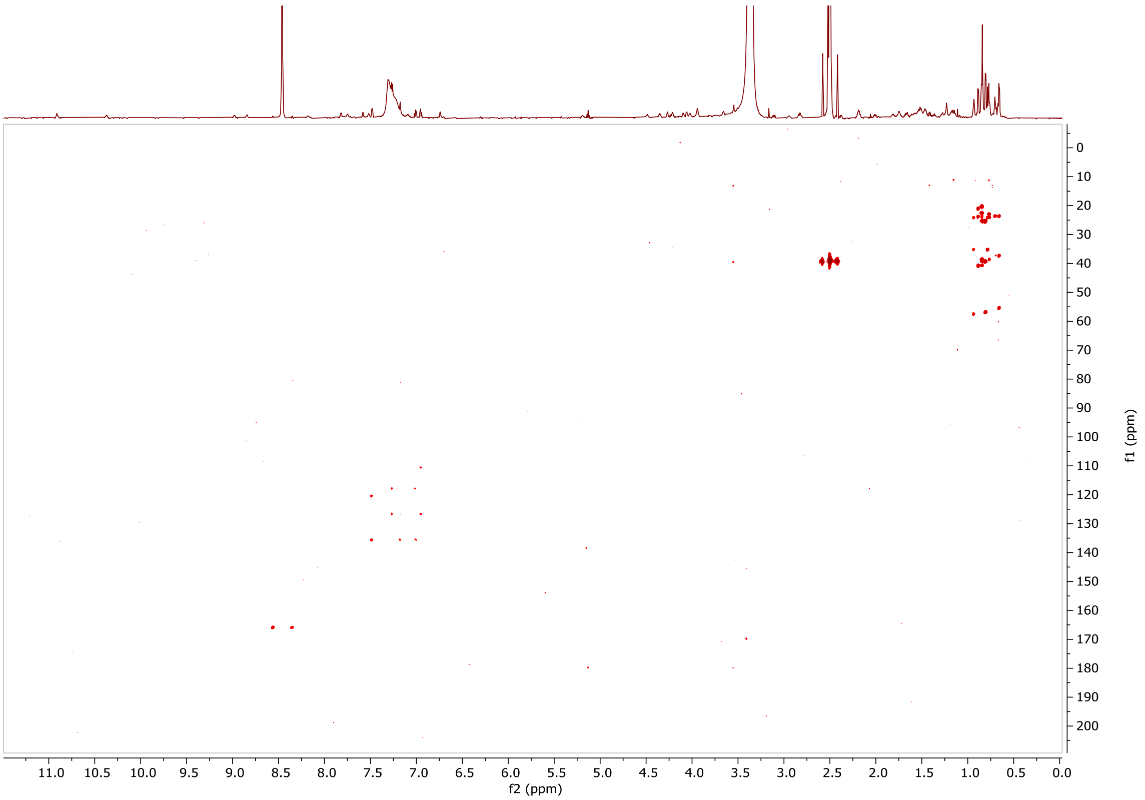


**Figure S15**. HBMC spectrum of **1** (850 MHz, in DMSO-$d_6$).

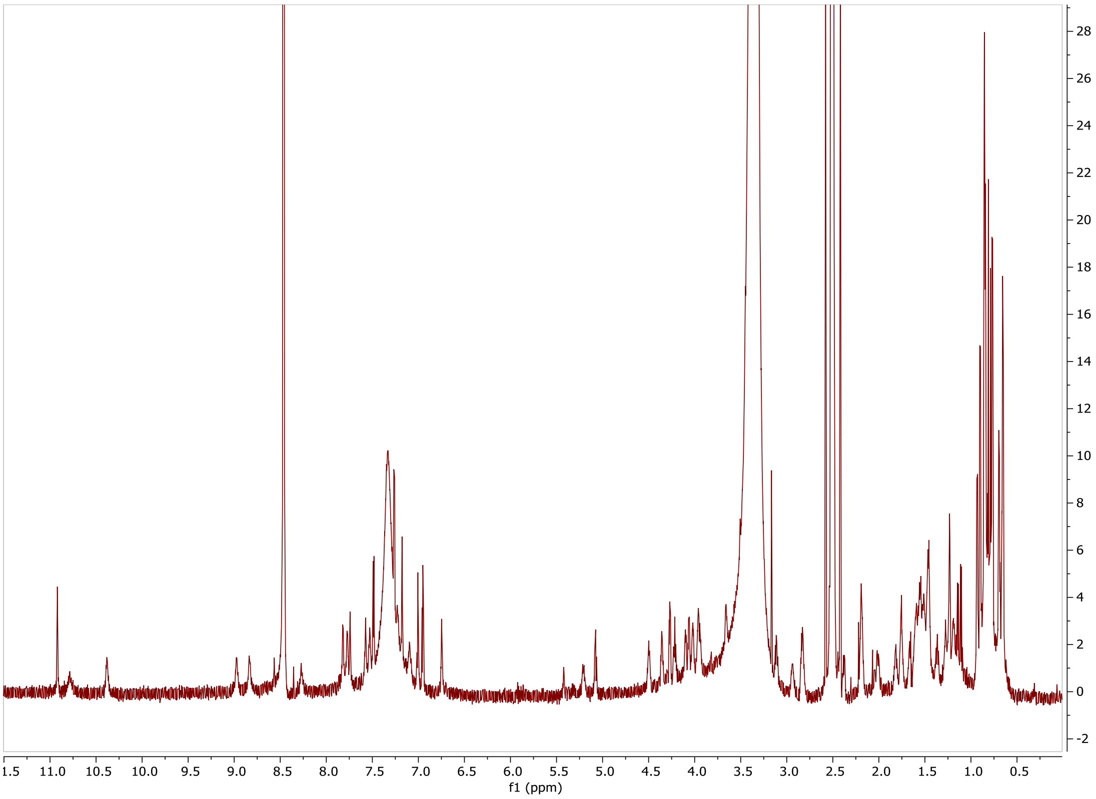


**Figure S16**. $^1$H NMR spectrum of **2** (850 MHz, in DMSO-$d_6$).

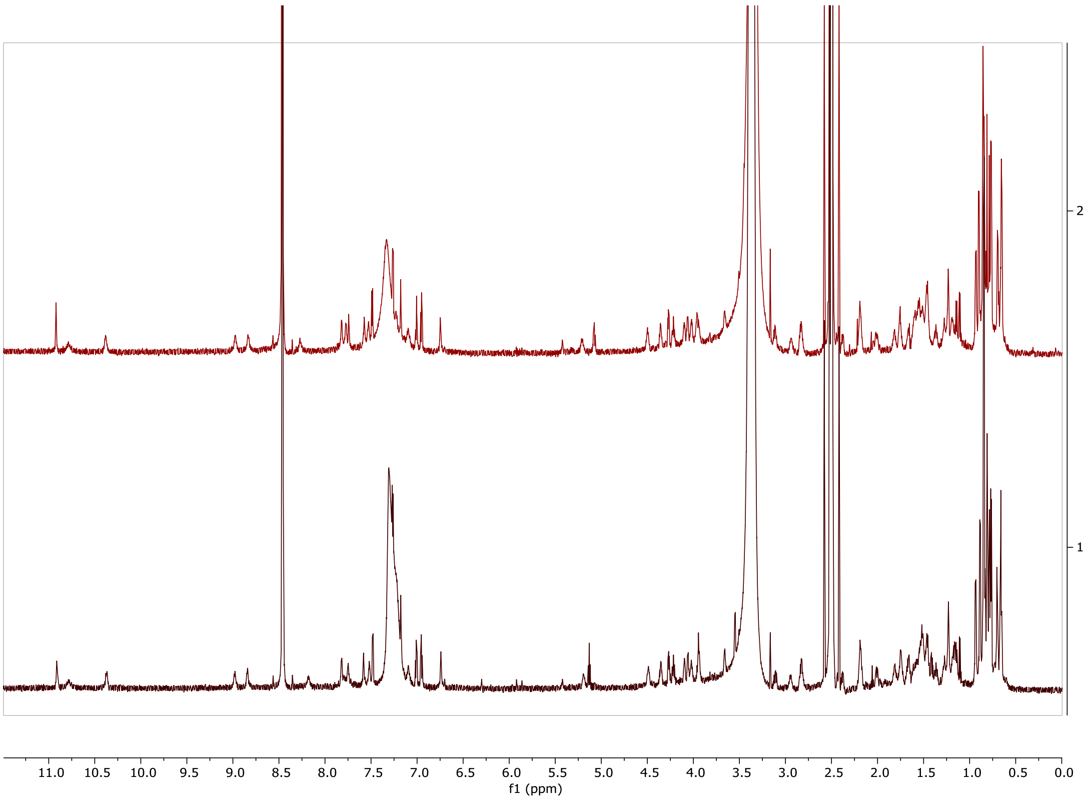


**Figure S17**. Stacked $^1$H NMR spectra of **1** and **2** (850 MHz, in DMSO-$d_6$).
